# Nucleotide-protocell interactions: A reciprocal relationship in prebiotically pertinent environments

**DOI:** 10.1101/2023.07.21.550025

**Authors:** Kshitij Deshpande, Sahil Sunil Mulewar, Sudha Rajamani

## Abstract

Spontaneous interactions between nucleotides and lipid membranes are likely to have played a prominent role in the emergence of life on Earth. However, the effect of nucleotides on the physicochemical properties of model protocellular membranes is relatively less understood. To this end, we aimed to discern the effect of canonical nucleotides on the properties of single-chain amphiphile membranes under prebiotically relevant conditions of multiple wet-dry cycles. Furthermore, the change in critical aggregation concentration of the membranes, and their stability in the presence of nucleotides, was also investigated in Astrobiologically analogue environments. We report that different nucleotides, lipid headgroups, and the ionic makeup of the system affect lipid-nucleotide interactions, which in turn can modulate the effect of nucleotides on the membranes. Specifically, oleic acid membranes seemingly showed an increase in the critical aggregation concentration, and an increase in the stability against prebiotically relevant selection pressures, in the presence of certain nucleotides. Our work has implications for understanding how nucleotides might have shaped the protocellular landscape of the prebiotic Earth.

## 1. Introduction

Compartmentalization is a primary attribute of cellular life, and hence, the emergence of compartments is considered a central step in the emergence of the earliest protocells.^[1–3]^ Life is thought to have originated in chemically heterogeneous environments, where different chemical entities would have spontaneously interacted with each other, affecting each other’s properties.^[4,5]^ Fatty acids and nucleotides are found abundantly in extant life and are also considered prebiotically relevant building blocks of protocells.^[6]^ Due to their importance in both extant biology and in the context of the earliest protocells, understanding how the interactions between these two molecular classes in the prebiotic milieu could have affected each other’s properties, and, in turn, their evolution, would have been significant for processes that led to the emergence of life. The effect of lipids on the properties of nucleotides has been well characterized in previous literature. Under wet-dry cycles, lipids have been shown to facilitate organization of nucleotides,^[7]^ which has been demonstrated to facilitate their oligomerization.^[8–10]^ Furthermore, the presence of lipids have shown to increase the stability of nucleotides in both the monomeric and oligomeric forms.^[10]^ Nucleotides are thought to interact with the membranes via salt-bridge interactions in the presence of divalent cations.^[11–13]^ However, in the absence of such cations, the interactions between nucleotides and membranes are thought to occur through hydrogen bonding, or hydrophobic interactions, especially in the case of fatty acid membranes.^[14–17]^ The interactions with nucleotides also affect some properties of lipid membranes. For example, oligonucleotides increase the permeability of phospholipid membranes in the presence of divalent cations.^[12,18]^ Nucleotide monophosphates (NMP) have been shown to affect the lamellarity and the lamellar thickness in phospholipid membranes due to surface localization.^[19]^ In a more prebiotically relevant context, the building blocks of nucleotides such as nucleobases and ribose, localize on the surface of decanoic acid membranes and increase their stability against higher concentrations of monovalent cations.^[14,15]^ There are noteworthy findings on the effect of nucleotides and its components (such as nucleobases and sugars) on the binding, stability and, to some extent, the self-assembly of membranes. Nonetheless, to the best of our knowledge, there are no studies that provide a comparative understanding of the effect of different nucleotides on the physicochemical properties of prebiotically relevant protocellular membranes. Additionally, we also wanted to study if interactions with nucleotides could confer any tangible “benefits” to the protocellular membranes, potentially affecting their fitness in a putative prebiotic landscape.

We investigated the effect of canonical ribonucleotides on fatty acid vesicles made entirely of oleic acid (OA) or a binary mixture of oleic acid and glyceryl monooleate (GMO). To investigate the effect of nucleotides on the physicochemical properties of the protocellular membranes, we used fluorescence spectroscopy tools that were recently developed by our group.^[20]^ Specifically, studies were carried out using fatty acid-based vesicles and disodiated salts of nucleotides, in buffered and in prebiotically analogous geothermal conditions. These experiments were conducted over multiple wet-dry cycles (unless mentioned otherwise) given their geological relevance on the early Earth, just as on extant Earth. ^[21]^ Using microscopy imaging with criteria that were previously standardized in our lab,^[22]^ we investigated the effect of nucleotides as co-solutes on the formation and stability of vesicles in prebiotically pertinent conditions. Our results demonstrate that nucleotides do affect the physicochemical properties of protocellular membranes such as micropolarity, membrane order and fluidity. We also show that the properties of the nucleobase (in the nucleotide) and the polar headgroup (of the lipid) affect nucleotide-membrane interactions, which in turn influence the properties of the membranes. Pertinently, experiments conducted in Astrobiologically relevant analogue conditions caused distinctive changes to the properties of the membranes, when compared to their buffered counterparts. Incidentally, the critical aggregation concentration of OA seemed to increase when compared to the control system in the presence of some of the nucleotides. Under Astrobiologically relevant analogue conditions, CMP seemed to increase the stability of OA as well as the binary OA-GMO vesicles, while GMP did not show such an effect. Also, OA system resulted in stable vesicle formation in the presence of all other nucleotides except for GMP in conditions of low buffer concentration and at low pH. Taken together, our results provide an exciting insight into nucleotide-membrane interactions and some of the factors affecting them.

## 2. Results and Discussion

In this study, we used vesicles that were made of oleic acid (OA) or a binary mixture of oleic acid and glyceryl-1-monooleate (GMO) in a 2:1 ratio. Disodiated salts of canonical ribonucleotides (5’NMP) were used as co-solutes. Relative concentrations of lipid and nucleotides were maintained at 1:5 in all the experiments. Lipids were suspended in 200 mM of pH 8.5 bicine buffer, or water samples from Astrobiologically relevant geothermal pools depending on the experimental conditions. The physicochemical properties and stability of the membranes interacting with the nucleotides (test systems), were compared over multiple wet-dry cycles with a control system containing 40 mM NaCl as a co-solute (termed Cont_N_). This was to eliminate any artefacts arising from changes in molality and excess Na^+^ counterions in the system as a consequence of the addition of NMP (See supplementary Information, Figure S4). To investigate the inherent effect of wet-dry cycles, the test systems were compared with model membranes in absence of co-solutes (control). Samples were taken after a set number of wet-dry cycles and subjected to further analysis.

### 2.1 Nucleotides influence the physicochemical properties of model protocell membranes under buffered conditions

The effect of nucleotide-membrane interactions on the physicochemical properties of single-chain amphiphile model protocell membrane systems was investigated. We characterized the micropolarity of the membranes, molecular order of lipids, fluidity of the bilayer and the turbidity of the resultant suspensions. These experiments were conducted in buffered conditions at pH 8.5. To gain a systems-level perspective of the effect of nucleotides on the membranes, the aforesaid properties were compared with the Cont_N_ over multiple wet-dry cycles. This was done using pyrene and nile red, which are hydrophobic fluorophores whose emission spectra are affected by the polarity of the microenvironment in which they are situated.^[23,24]^ The emission spectrum of pyrene shows five vibronic peaks between the wavelengths of 370 nm and 400 nm (Figure S2). The ratio of intensities between the 1^st^ peak representing the 0-0 band transition (I1) at 372 nm, and the third peak at wavelength 383 nm (I3), i.e., I_1_/I_3_, has been shown to be directly correlated to the polarity of pyrene’s microenvironment.^[20,23]^ On an average, the I_1_/I_3_ ratio for the membranes interacting with pyrimidines was higher than the ones interacting with purines (Figure 1a). Over the wet-dry cycles, the change in the micropolarity was statistically significant for membranes interacting with GMP and CMP (Table 1). For systems interacting with CMP in this context, the micropolarity increased, while it decreased in the case of interaction with GMP. This decrease in the micropolarity in the test systems containing GMP in the vicinity could potentially be due to the efficient stacking and self-base-pairing of GMP at concentrations of 20 mM.^[25,26]^ This could, in principle, lead to the exclusion of water from the midst of GMP molecules that are interacting with the membrane, thereby decreasing the polarity of the membrane. Pyrimidines, on the other hand, do not stack as efficiently. Hence, their spread is likely to be more uniform across the surface of the bilayer compared to the purines; potentially decreasing the packing order and in turn increasing the water accessibility.

**Figure 1.**
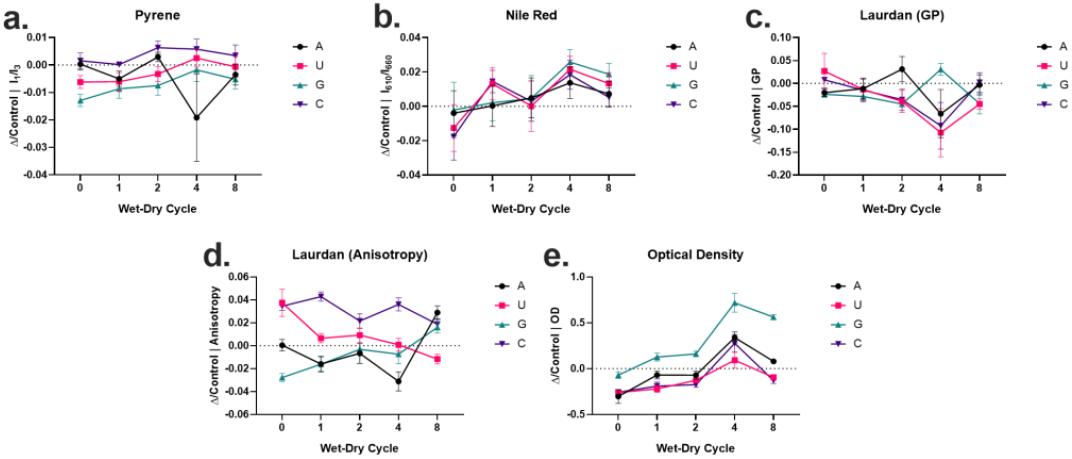
Nucleotides affect the physicochemical properties of homogenous OA membranes in laboratory buffered conditions. The Y-axis of the scatter plots represents normalized differences in properties calculated against Cont_N_ for a) I_1_/I_3_ of pyrene, b) I_610_/I_660_ of nile red, c) GP of laurdan, d) fluorescence anisotropy of laurdan, e) optical density. Membranes were prepared in 200 mM Bicine buffer at pH 8.5 and Cont_N_ is the control sample of pure OA with NaCl as co-solute. Readings were compared over wet-dry cycles; the timepoints have been appropriately indicated at the x-axis of each data set (as 0-8), and the nucleotide type has been indicated on the right. N = 4, error bars = std. error.

**Table 1.** Comparison of p-values obtained from two-way Repeated Measures ANOVA analysis performed for the test samples and the Cont_N_ for homogenous OA vesicles. The physicochemical properties were compared for the test systems with Cont_N_ over eight wet-dry cycles to gain a systems level perspective of the effect of nucleotides on the physicochemical properties of protocellular membranes. p-values smaller than 0.05 were considered statistically significant and highlighted in bold text.

| Co-solute | AMP | UMP | GMP | CMP |
| --- | --- | --- | --- | --- |
| Membrane Property |  |  |  |  |
| **Micropolarity near the headgroup** | 0.2388 | 0.1102 | **0.0025** | **0.0424** |
| **Micropolarity at the center of bilayer** | 0.5451 | 0.3976 | 0.2089 | 0.5496 |
| **Molecular order of lipids** | 0.1882 | **0.0028** | **0.0156** | **0.0354** |
| **Membrane fluidity** | 0.1274 | **0.0403** | 0.0645 | **<0.0001** |
| **Turbidity** | 0.1732 | **0.0004** | **<0.0001** | **0.0039** |

Along with pyrene, we investigated the effect of nucleotides on the polarity of the model membranes using fluorescence emission properties of nile red. The spectrum of nile red shows a blue shift in its emission maximum in non-polar environments. For pure oleic acid membranes, the emission maximum was observed at 626 nm. Earlier literature has shown that the ratio of emission intensities at wavelengths 610 nm (I610) and 660 nm (I660) is inversely proportional to the polarity of nile red’s microenvironment.^[20,27]^ The I_610_/I_660_ of nile red did not show any statistically significant change in any of the test systems when compared with the Cont_N_ system (Figure 1b). The difference in the micropolarity detected by pyrene and nile red is likely a consequence of their different localization patterns across the lipid bilayer, where pyrene is thought to situate itself closer to the polar headgroup,^[28]^ while nile red is present towards the center of the bilayer.^[29]^

Since lipid-nucleotide interactions are likely to mainly be surface interactions,^[15]^ they are also poised to impart new phase properties on the model protocell membranes. We studied the structural order of lipids and the fluidity of the bilayer using fluorescence emission properties of laurdan whose steady-state emission allows for this to be done effectively. A parameter called generalized polarization (GP) can be obtained, which is defined based on the steady-state emission intensities of laurdan at wavelengths 430 nm and 500 nm (see methods section). The GP is directly correlated with the structural order of lipids in the membrane.^[30,31]^ GP is dependent on the motion of water molecules in the vicinity of laurdan, hence it is also correlated with water accessibility around laurdan molecules.^[32]^ On an average, we observed a decrease in the GP value in the test systems, indicating increased water activity in the bilayer. This also indicated a decrease in the structural order in the oleic acid membranes (Figure 1c). The decrease in order, however, was statistically significant for test systems interacting with UMP, GMP, and CMP, but not the ones interacting with AMP (Table 1). Laurdan shows anisotropy in its fluorescence intensity, which can also be used to study the fluidity of the bilayer. The fluidity of the bilayer is related to the motion of lipid molecules in the bilayer. This in turn affects the anisotropy of laurdan, which is inversely correlated with the fluidity of the membrane.^[32]^ When compared with Cont_N_, fluorescence anisotropy showed an overall increase for the test systems containing pyrimidines as co-solutes and a minor decrease for the systems containing purines (Figure 1d). The changes in the membrane fluidity over multiple wet-dry cycles were statistically significant only for test systems containing pyrimidines. In the presence of pyrimidines, the membrane seemed to have become comparatively rigid. Here, we observe an inverse correlation between the changes in structural order and the fluidity of the bilayer. This is in accordance with earlier reported studies where the surface interactions with small molecules have shown to result in independent changes in the membrane order and fluidity.^[33,34]^ This indicates the existence of a spatial component in the effect of nucleotide-membrane interactions on the physicochemical properties on the model membranes, whereby the nucleotides have varying effect on the membranes across the bilayer.

Next, we checked the turbidity of the model lipid membranes, which provide a general information about different properties of the vesicles, including their average size, lamellarity, and bilayer thickness.^[35]^ Turbidity was analyzed using optical density (OD) measurements in a UV-Visible spectrophotometer at a wavelength of 450 nm. The interactions with nucleotides resulted in a net decrease in the OD for all the test samples (except GMP) compared to Cont_N_ (Figure 1e). This change was statistically significant in the case of the aforementioned test systems when compared with Cont_N_, over multiple wet-dry cycles (Table 1), indicating a possible change in the aforesaid bulk properties of these membranes. This seems to indicate that the properties of the nucleobases (of the nucleotides) affect the nucleotide-membrane interactions, in turn affecting the physicochemical properties of the model membranes.

Since nucleotides are thought to interact with membranes mainly through surface interactions, we hypothesized that the properties of the headgroup of the lipids are also likely to play an important role in these interactions. Since fatty acids form vesicles through pseudo-di-acyl based interactions,^[36]^ its net hydrogen bonding character is likely to be as an acceptor.^[14,15]^ This potentially would result in the interactions with nucleotides via hydrogen bonds with the nucleobase and the sugar moieties of the nucleotide.^[15]^ The inclusion of a hydrogen bond donor in the bilayer could, therefore, result in additional interactions with the nucleotides, potentially changing the manner in which they affect the membranes. To investigate this, we used model protocell membranes that were prepared using a binary mixture of OA and glyceryl-1 monooleate (GMO) in a 2:1 ratio. This is because the GMO has hydride groups that can potentially act both as hydrogen bond donors and acceptors. Previous studies also showed that OA-GMO binary membranes are more stable compared to pure OA membranes against prebiotically relevant selection pressures, highlighting their relevance in the prebiotic world.^[37]^

We then conducted similar experiments with the binary membranes as with the pure OA membranes (as described in the previous sections). The micropolarity was affected for the binary lipid membrane system that was observed in studies undertaken with both pyrene and nile red. However, the nucleotides that affected the micropolarity were distinct from the pure OA membranes when compared with their respective Cont_N_ (Table 1, Table ST1). Interestingly, we observed contrasting changes in the micropolarity for pyrene and nile red. For the membranes interacting with AMP, pyrene showed a net decrease in the polarity while the nile red showed a net increase (Figure S5a, S5b). This is likely a consequence of the spatial nature of the nucleotide-membrane interactions. The laurdan GP did not significantly change in the binary membrane systems, however, the anisotropy showed a net decrease compared to the Cont_N_ across the test systems studied (Figure S5c, S5d, Table ST1). OD showed an overall decrease in the test systems interacting with purines, whereas there was an increase in the systems interacting with pyrimidines (Figure S5e), and the change was statistically significant for all the test systems as against the Cont_N_. We suspect the deviation in changes for the OA-GMO binary system from the ones that were observed in the pure OA system, is likely a consequence of the comparatively higher hydrogen bond donating character of the OA-GMO binary membranes. These are likely to influence the nature of interactions with the nucleotides. Hence, the ability of the lipid headgroups to form non-covalent interactions also considerably affects the influence of nucleotides on the physicochemical properties of the model membranes.

### 2.2 Nucleotides affect critical aggregation concentration of oleic acid membranes

In order to understand the effect of nucleotide-membrane interactions on the formation of model protocellular membranes, we studied the critical aggregation concentration (CAC) of OA. We calculated the CAC of OA in the presence of different nucleotides using diphenyl hexatriene (DPH) fluorescence intensity-based assay developed in our lab earlier.^[37]^ DPH is a fluorescent dye that shows a substantial increase in the emission intensity in non-polar environments, such as the hydrophobic tail region of self-assembled structures of OA (when compared to the aqueous medium). Fluorescence intensity was monitored at various concentrations of OA to measure the CAC (see methods section). The fluorescence intensity was plotted against the lipid concentration, and fitted to equation 1 to find the *x*°, which is the point of inflection in the fluorescence intensity that represents the CAC.

The CAC of the OA control system (without any co-solutes) was found to be around 0.042 mM (Figure 2a). Addition of NaCl (Cont_N_) showed no significant change in the CAC (Figure 2b). The test sample with AMP as co-solute also did not show any significant change in the critical aggregation concentration (Figure 2c). However, the other nucleotides significantly increased the CAC (Figure 2d-f). The increase in the CAC in the presence of CMP, GMP and UMP, as reported by the DPH fluorescence assay, indicates that nucleotides likely affect the formation of the self-assembled structures of OA. However, further investigation is required to explain how some nucleotides could be affecting the critical aggregation concentration.

**Figure 2.**
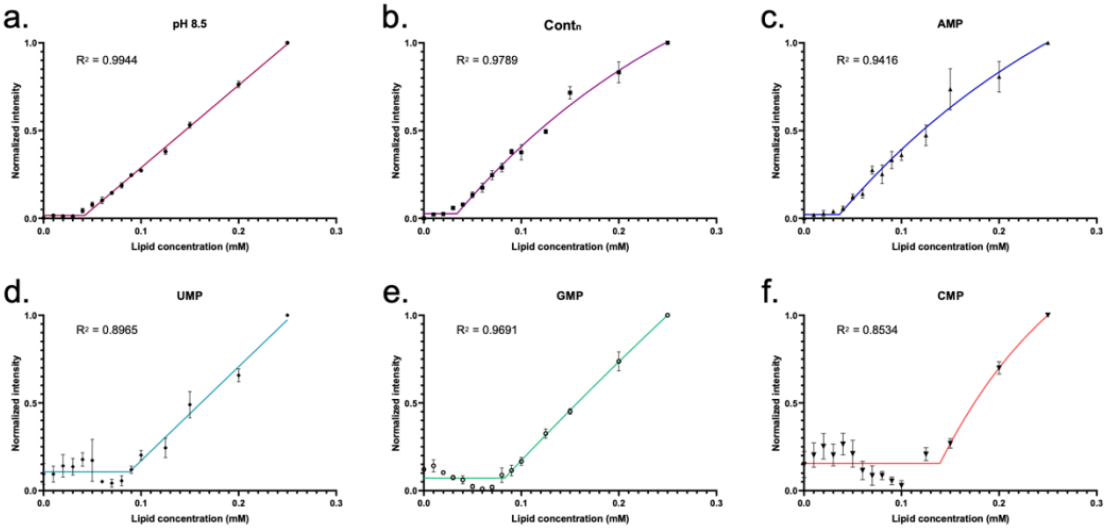
Critical aggregation concentration of oleic acid is affected by the presence of ribonucleotides as co-solutes. Normalized fluorescence intensity of DPH plotted against concentrations of oleic acid for a) OA control, b) OA + NaCl (Cont_N_), c) OA + AMP, d) OA + UMP, e) OA + CMP, f) OA + GMP. The dotted line represents the critical aggregation concentration calculated for the given system as elaborated in the methods section. CAC for suspension a) Control = 0.042 ± 0.002 mM, b) Cont_N_ = 0.034 ± 0.004 mM, c) AMP = 0.036 ± 0.006 mM, d) UMP = 0.088 ± 0.008, e) GMP = 0.083 ± 0.005, and f) CMP = 0.139 ± 0.009. N = 3, error bars = std error.

### 2.3 Effect of wet-dry cycles on model protocellular membranes

Wet-dry cycles are considered crucial for the oligomerization of nucleotides,^[8]^ and have also been shown to affect some of the properties of model membranes,^[20]^. Hence, we investigated how the properties of a particular system changes as a consequence of the wet-dry cycles. These effects were studied for a given membrane system by comparing the values for a property e.g., I_1_/I_3_ for pyrene, before starting the wet-dry cycles as against values after completing the N^th^ cycle, using a two-tailed t-test (see methods section). Amongst the properties studied, only laurdan fluorescence anisotropy and OD were significantly affected by the wet-dry cycles for both the pure OA (Figure 3a, Table ST3) and the OA-GMO binary membrane systems (Figure 3b, Table ST4). On the other hand, the micropolarity and the structural order of these membranes were unaffected for the control systems as well as the test systems (Figure S6, Table ST3, ST4).

**Figure 3.**
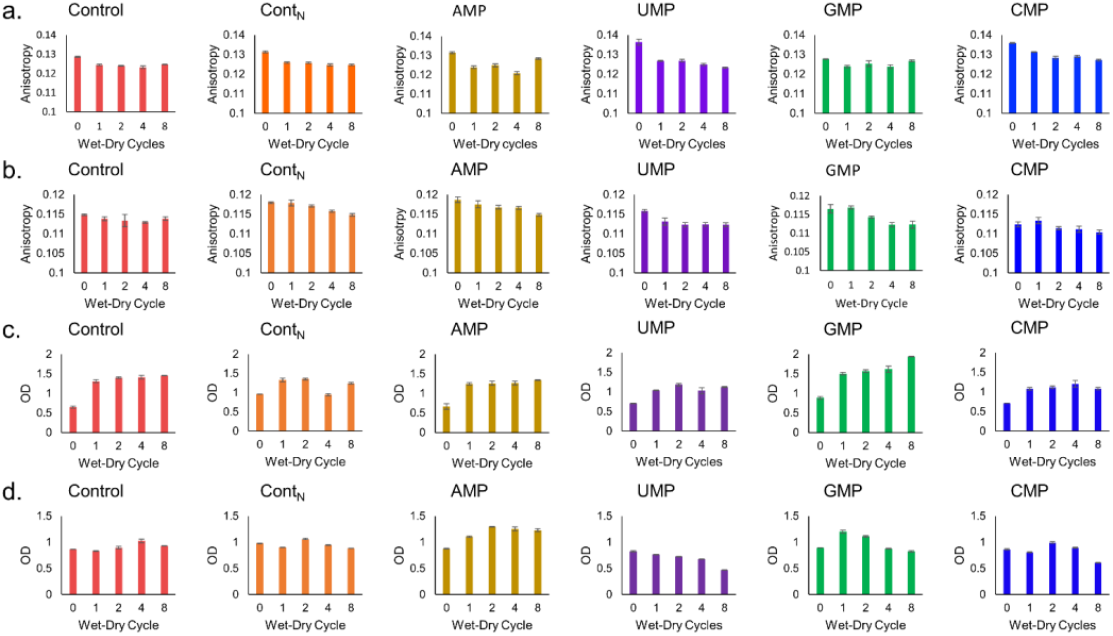
The wet-dry cycles, in laboratory buffered conditions, significantly affect a) fluorescence anisotropy for OA systems, b) fluorescence anisotropy of OA-GMO binary systems, c) optical density for OA systems, d) optical density for OA-GMO (2:1) binary systems. Membranes prepared in 200 mM Bicine buffer at pH 8.5. Timepoints have been appropriately indicated on the x-axis of each data set (as 0-8). The wet-dry cycles were conducted at 90°C under a gentle flow of N_2_. N=4, error bars = std error.

Laurdan fluorescence anisotropy showed a consistent decrease over the wet-dry cycles for both the pure OA and the OA-GMO binary membrane systems (Figure 3a, 3b). This decrease in the laurdan fluorescence anisotropy could potentially stem from a net decrease in the average vesicle radius over the wet-dry cycles, as noted by Sarkar et al. previously.^[20]^ To check if this were the case, we compared the fluorescence anisotropy of pure OA membranes prepared using only thin-film rehydration with a reported average radius of ∼500 nm^[20]^ with extruded membranes with an average radius of ∼50 nm. For oleic acid membranes both in the absence of any co-solutes and with AMP, the extruded vesicles showed a statistically significant increase in anisotropy compared to the non-extruded ones, disproving the possibility that the decrease in membrane anisotropy was a consequence of decreasing membrane radius over the wet-dry cycles (Figure S7). The fluidity of the bilayers seemed to increase with the wet-dry cycles, and this increase did not seem to be related to the average membrane size.

Turbidity was significantly affected in both the pure OA membranes as well as the OA-GMO binary membranes. The OD of the OA membranes increased after the first wet-dry cycle and plateaued thereafter (Figure 3, Table ST3). OA-GMO membranes did not seem to follow any particular pattern across the control and test systems (Figure 3, Table ST4). Since turbidity is representative of multiple properties of the membranes, they are likely affected due to wet-dry cycles. However, further experimental and theoretical studies are required to understand the exact mechanism through which wet-dry cycles affect the membranes both in general and during the wet-dry cycling process.

### 2.4 Nucleotides influence the physicochemical properties of model protocell membranes under Astrobiologically relevant conditions

Our results so far were obtained under controlled laboratory buffered conditions. Even though such experiments provide valuable insights into the interactions of nucleotides with the membranes, they fail to account for the complexity of natural geochemical settings where life is thought to have originated. Hence, investigating these interactions in Astrobiologically analogous environments would provide more realistic understanding of the interactions that might have panned out on the early Earth. Terrestrial geothermal pools are one of the prominent geological niches where life is thought to have originated.^[4,21,38]^ Geothermal pool-like environments are known to facilitate prebiotically relevant events including the assembly of single-chain amphiphiles into compartments, considered crucial for the emergence of the earliest protocells.^[38]^ Therefore, we simulated these conditions using water samples collected from an Astrobiological analogue geothermal pool located in Ladakh, India.^[39]^ The water samples from Puga contain a variety of different cations and anions. As nucleotide-membrane interactions are also dependent on the ionic species in the system, especially cations, the ionic diversity is likely to affect the interaction between the two entities. The effect of nucleotide interactions was studied with the OA-GMO binary membrane system that was subjected to multiple wet-dry cycles using Puga water (see methods section). The binary membranes are stable in the Puga water; on the other hand, pure OA membranes were found to be unstable under these conditions.^[22]^

We compared the physicochemical properties of the lipid suspensions prepared in the analogue water samples with the ones prepared in bicine buffer, over multiple wet-dry cycles. Equation 3’, a modified version of the earlier equation (equation 3), was used to calculate the normalized difference between the values for the properties in the analogue conditions against the values in the buffered one (see methods for more details).

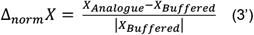

Pyrene showed an increase in the I_1_/I_3_ ratio, implying an increase in polarity in all the suspensions prepared using the analogue site water sample when compared to their buffered counterparts (Figure 4a). On the other hand, nile red showed a significant change in the polarity only for the GMP-based system when compared to its buffered counterpart (Figure 4b, Table ST2). The molecular order (laurdan GP) was not significantly affected in the analogue water sample while the membrane fluidity decreased for the Cont_N_ system but increased in the case of the test systems (Figure 4c, 4d). Similar to the fluidity, the Cont_N_ system showed an overall decrease in turbidity through the wet-dry cycles, while the test systems showed an increase when compared to their buffered counterparts (Figure 4e). It is pertinent to note that the water sample from Puga had a much lower net salt concentration (<30mM)^[22]^ when compared to the buffered systems, where the salt concentration was 200 mM. The lower salt concentrations, along with the wider variety of cations such as K^+^, Li^+^ and Ca^2+^ that are present in Puga, likely affected the interactions of nucleotides with the membranes, altering the effect they had on the OA-GMO binary membrane system in this process. Relevantly, these results allow us to examine the interactions between nucleotides and membranes in a more realistic setting that better resembles the conditions of the early Earth. Therefore, it is worthwhile to note that not only the properties of the nucleobases (of the nucleotides) and the polar headgroups (of lipids) are pertinent, but the ionic makeup of the suspension as well would have affected the physicochemical properties of the model protocellular membranes during their emergence and early evolution.

**Figure 4.**
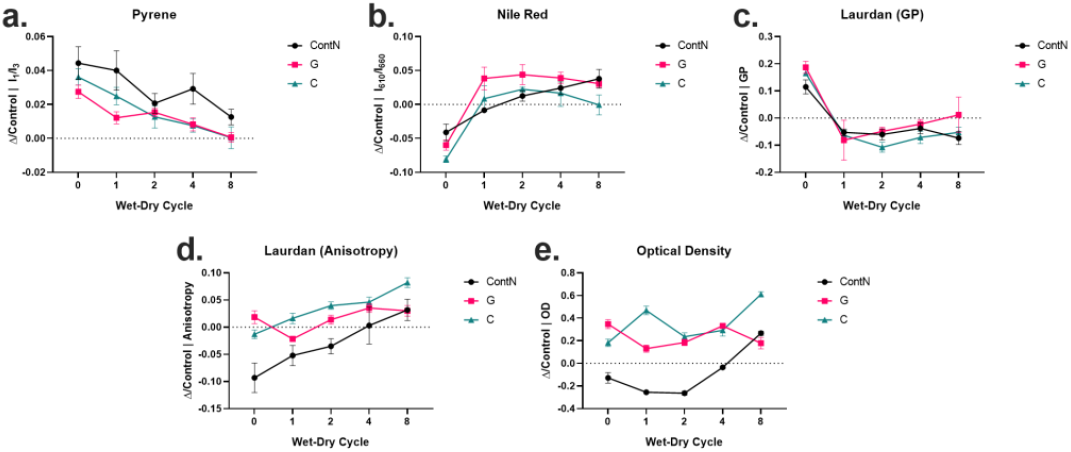
Astrobiologically relevant analogue conditions change the effect of nucleotides on the properties of membranes prepared with OA-GMO. Protocells were prepared in analogue conditions using Puga hot spring water sample and the properties were compared against membranes that were prepared in 200 mM bicine at pH 8.5. Normalized differences between the analogue and laboratory buffered conditions were calculated to understand the effect of analogue environment on the properties of protocellular membranes over wet-dry cycles for, a) I_1_/I_3_ of pyrene, b) I_610_/I_660_ of nile red, c) generalized polarization for laurdan, d) fluorescence anisotropy of laurdan and e) optical density. Systems legend has been indicated on the right side of the graphs. Experiments performed in independent experimental replicates. N = 4, error bars = std. error.

### 2.5 Effect of nucleotides on the stability of membranes against Astrobiologically relevant stresses

We investigated the effect of nucleotides on the stability of the SCA membranes under Astrobiologically relevant analogue conditions using water samples from terrestrial geothermal pools. Towards this end, we subjected both pure OA and the OA-GMO binary membrane systems to multiple wet-dry cycles in water samples collected from three different geothermal pools in Ladakh, namely Puga (PU), Chumatang (CH), and Panamic (PA). The net concentration of lipids was maintained at 4 mM across all the reactions. Water samples from these three pools have been shown to inhibit the formation of pure oleic acid vesicles, whereas the binary membranes of OA-GMO have been shown to readily form vesicles in them.^[22]^

The vesicle suspensions from our wet-dry cycled reaction samples were observed using differential interference contrast (DIC) microscopy to characterize different self-assembled structures in the system (see methods section). As expected, in absence of nucleotides, we did not observe any vesicles in pure OA suspensions in any of the analogue water samples (Figure 5, S8, S9, Table 2). As previously reported, in the suspensions containing OA-GMO in 2:1 ratio, lipids readily assembled into vesicles in water samples from Puga and Chumatang, but not in Panamic (Figure S8, S9, Table 2). Results obtained in these control experiments were consistent with those reported by Joshi et al.^[22]^ However, to our surprise, the test samples containing 20 mM CMP showed robust formation of vesicles in Puga for pure OA membranes in all the wet-dry cycles (Figure 5b). However, in Chumatang and Panamic, we observed vesicles only in isolated cycles and not throughout (Figure 5c, 5d). In the case of the binary membrane system, when CMP was added as a co-solute, we observed the formation of vesicles in water samples from all three geothermal pools (Figure S11). However, this increased stability was not observed when GMP was added as the co-solute (Figure S10). When GMP was added as a co-solute instead of CMP, we did not observe any vesicles for pure OA throughout the wet-dry cycles in any of the analogue water samples (Figure 11, Table 2). And, for the OA-GMO binary system, even after the addition of GMP, the vesicle formation was observed to be the same as the control system with vesicle formation observed only in Puga and Chumatang water samples. (Figure S12). Hence, the addition of CMP as a co-solute seemed to increase the stability of both pure OA and the binary OA-GMO membranes, however, GMP did not show any such phenomenon.

**Figure 5.**
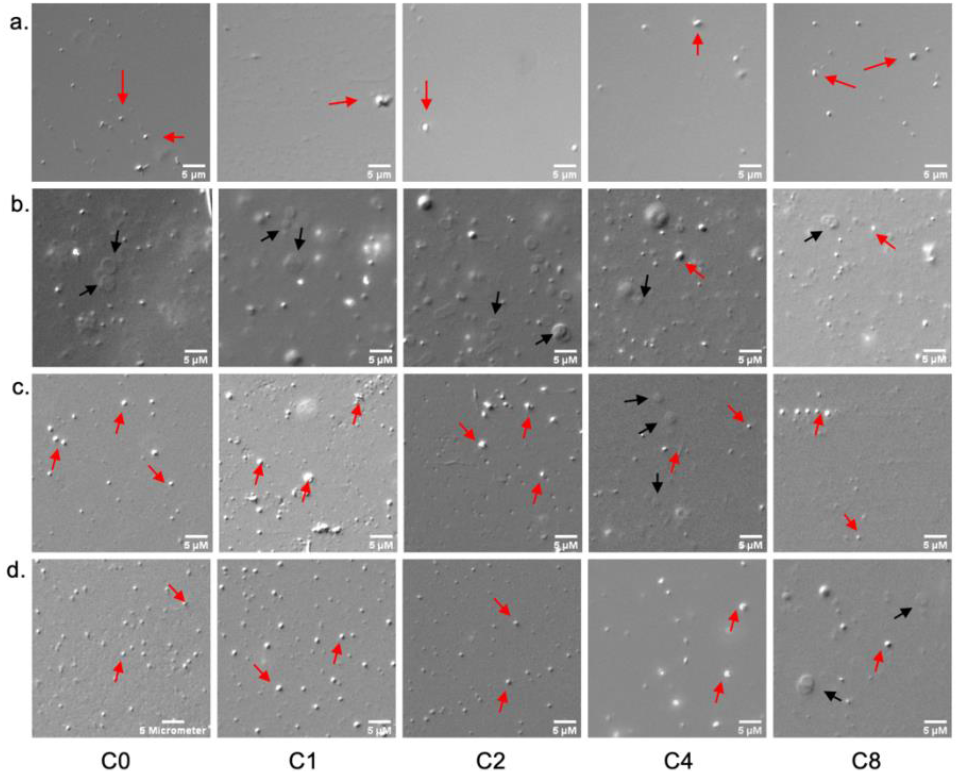
CMP increases the stability of OA membranes under Astrobiologically relevant analogue environments. a) Puga control (without co-solutes) b) Puga + CMP, c) Chumatang + CMP, d) Panamic + CMP. We observe oleic acid vesicle formation for all the suspensions prepared in water samples from geothermal pools. Timepoints have been indicated in the bottom panel as C0-C8. Samples were observed using DIC microscopy at 40x, scale bar is 5 μm, N=3. Black arrow= vesicles, blue arrow= oil droplets.

**Table 2.**
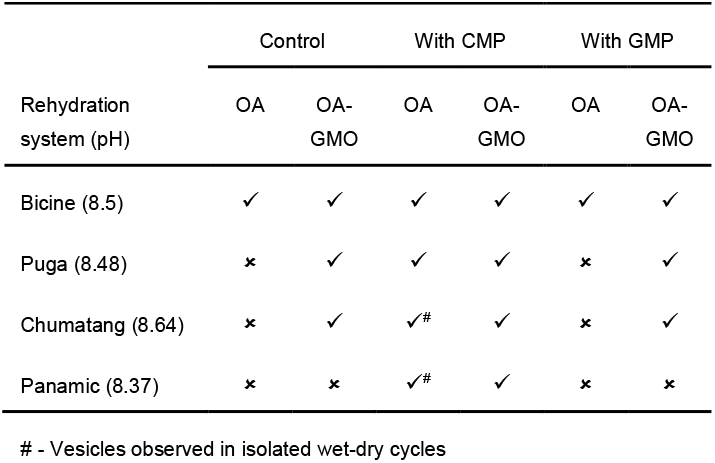
Results for vesicle formation experiments under buffered and prebiotically relevant analogue conditions, and the effect of nucleotides on vesicle formation. Buffered samples were prepared in 200 mM bicine at pH 8.5. Prebiotic analogue conditions were simulated using water samples from terrestrial geothermal pools of Puga, Chumatang and Panamic. The initial pH (without addition of lipids or nucleotides) of these systems is mentioned in parentheses. The pH for water samples from analogue geothermal pools was measured by Joshi et al..^[22]^ ✓ = Vesicles observed, ✗ = no vesicles observed.

|  | Control |  | With CMP |  | With GMP |  |
| --- | --- | --- | --- | --- | --- | --- |
|  | OA | OA-GMO | OA | OA-GMO | OA | OA-GMO |
| Rehydration system (pH) |  |  |  |  |  |  |
| Bicine (8.5) | ✓ | ✓ | ✓ | ✓ | ✓ | ✓ |
| Puga (8.48) | ✗ | ✓ | ✓ | ✓ | ✗ | ✓ |
| Chumatang (8.64) | ✗ | ✓ | ✓<sup>#</sup> | ✓ | ✗ | ✓ |
| Panamic (8.37) | ✗ | ✗ | ✓<sup>#</sup> | ✓ | ✗ | ✗ |

Intrigued by the effect of CMP on the stability of the OA membranes, we investigated the stability of pure OA to form vesicles with canonical nucleotides (AMP, UMP, GMP, CMP) in varying buffer concentrations without wet-dry cycles to check for the effect of nucleotides against lower salt concentration and pH in the suspensions. A 200 mM stock of bicine buffer set at pH 8.5 was diluted to 0 mM, 5 mM, 10 mM, 20 mM, 50 mM, 75 mM and 100 mM using an appropriate amount of Milli-Q. The pH of the buffer dilutions was not adjusted after dilution. When suspended in these buffers, addition of OA did not change the pH of the buffers except when it was suspended in pure Milli-Q (Table ST5). After the addition of nucleotides, the pH was restored to 7 in the system suspended with Milli-Q. The control system did not show the formation of vesicles below the bicine concentration of 50 mM (pH <8.5). Interestingly, we observed stable vesicles in all the test systems except GMP-based ones over a larger range of salt concentrations and pH (Figure 6, Figure S13). The increased stability seemingly conferred by the nucleotides (except GMP) shows that nucleotides likely stabilize model protocellular membranes against lower salt concentrations. Since OA vesicles are most stable at a pH of 8.5,^[37]^ stability was also seemingly conferred against lower pH as well.

**Figure 6.**
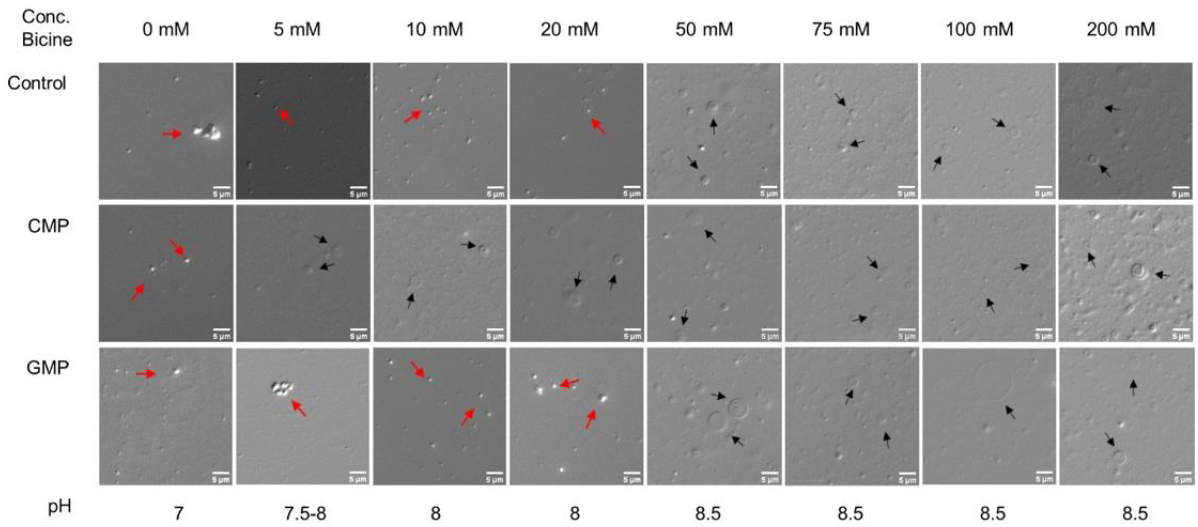
Nucleotides increase the stability of oleic acid vesicles against buffer dilution and pH changes. Buffer concentration has been indicated in the top panel, while the pH is indicated in the bottom panel. Control reaction and the reactions containing GMP form vesicles only at pH 8.5 and at buffer concentrations of 50 mM and above. However, suspensions containing CMP form vesicles even when the buffer concentration is down to only 5 mM at a pH range between 7.5-8. Samples were observed using DIC microscopy at 40x, and the scale bar is 5 μm, N=2. Black arrow= vesicles, red arrow= oil droplets. For overall comparison of different systems refer Table ST6.

The inability of only GMP to stabilize OA vesicles rules out the possibility of the Na^+^ counterions of nucleotides being the causative agent in the process. The lack of pH changes in the buffer dilution experiments rules out the possibility that the nucleotides conferred stability by changing the pH of the suspension. The orientation of nucleotides interacting with the membranes is likely a key factor affecting membrane stability. As for the curious case of GMP, at concentrations of 20 mM it is known to efficiently form self-interactions such as pi-interactions, and self-base pairing.^[25,26]^ These potentially could make GMP unavailable to interact with the membranes, or result in larger domains of GMP on the surface of a membrane. These interactions could result in the effect of GMP on the surface of the membrane being localized only to particular regions, and not in a uniform manner as might be the case with the other nucleotides (Figure S14). However, more experimentation is required to ascertain the mechanistic underpinnings behind the likely increase in the stability of the membranes against Astrobiologically pertinent stresses due to nucleotide-membrane interactions.

## 3. Conclusion

The spontaneous interactions between nucleotides and lipid membranes lead to a variety of emergent phenomena that are likely to have significant implications on each other’s properties in the prebiotic milieu. Even though the effect of lipids on the properties of the nucleotides has been well established over the years, the other side of these interactions, i.e., the effect of nucleotides on model protocellular membranes has remained relatively less explored. Our goal in this regard was to understand what the presence of different nucleotides meant for the physicochemical properties of the studied model membranes. After admixing disodiated salts of canonical nucleotides, the model protocellular membranes were subjected to multiple wet-dry cycles to investigate the effect of nucleotide-membrane interactions over multiple assembly-disassembly cycles. The effect of different nucleotides and lipid headgroups on the nucleotide-membrane interactions was evaluated, some of which affected the membranes in both buffered and Astrobiologically relevant analogue conditions. We also investigated the potential effects of nucleotides on the formation and stability of the membranes against Astrobiologically relevant stresses.

As the prebiotic milieu is thought to be a chemically heterogenous system, it is reasonable to assume that the local environment would have played an important role in affecting the properties of the membranes. Co-solutes and environmental factors such as wet-dry cycles would have facilitated a variety of interactions for the protocellular membranes by acting as prebiotically relevant selection pressures that would have impinged on their sustenance and evolution. Learning the effect of these emergent phenomena and the selection pressures is fundamental to understanding how extant life evolved the way it did. Earlier literature, including work from our lab, has investigated the nature of the self-assembly of single-chain amphiphile membranes in prebiotically relevant contexts.^[16,20,22,37,40]^ In this study, we undertook a comparative analysis of the effect of different canonical nucleotides on the self-assembly of model protocellular membranes, with the intent of gaining an understanding of the relevance of such interactions in the context of the early Earth.

Our results demonstrate that the non-covalent interactions of membranes with nucleotides could significantly affect the properties of model protocellular membranes. The wet-dry cycles affected the turbidity and the fluidity of the model membranes; however, the effect of wet-dry cycles was relatively consistent across the control and test systems. The interactions with nucleotides caused changes in the physicochemical properties of the membranes such as the micropolarity, order and fluidity of the model membranes. In all, our results demonstrate that different nucleotides, based on the properties of the nucleobase, affect the membranes in a different manner, and these interactions likely have a spatial component to them across the bilayer. Just like the nucleobases, the properties of the lipid headgroups also affect nucleotide-membrane interactions. The effect of nucleotides on the OA-GMO binary membranes differed from what was observed with the pure OA membranes. We attribute this difference to the higher hydrogen bond donating character of the GMO moiety in the binary membrane systems. Along with the properties of the nucleobases and the lipid headgroups, the inorganic ionic species in the suspension also affected the properties of the membranes. When the OA-GMO binary membranes were subjected to analogue conditions, the micropolarity, fluidity and turbidity seemed to be significantly affected compared to their buffered counterparts.

Our results also highlight the effect of the nucleotides on the formation and stability of the membranes. The critical aggregation concentration of oleic acid was significantly affected by the presence of CMP, GMP and UMP. The effect of nucleotides on the CAC when compared with the Cont_N_ showed that the nucleotides somehow affected the formation of vesicles, but did however stabilize the already formed vesicles when they were present in concentrations above the critical aggregation concentration. Some of them also seemed to increase the stability of the membranes in Astrobiologically relevant conditions involving geothermal pool water samples, as well as against lower salt concentrations and relatively lower pH conditions.

In conclusion, this study shows that nucleotides as co-solutes do affect the self-assembly properties of the single-chain amphiphile membranes. Similarly, nucleotides also affect their stability. Pre-existing literature has highlighted the role of nucleotide-membrane interactions from the perspective of lipids where it was shown that lipids conferred discernible benefits to the rate and extent of oligomerization,^[8]^ as well as the stability of the nucleotides and their oligomers.^[10]^ In this work, we provide evidence for nucleotides conferring tangible benefits to the lipid membranes. Hence, in the context of previously reported lipid-nucleotide literature, we establish a reciprocal relationship between nucleotides and membranes with reasonable confidence (Figure S15). Their co-existence would have influenced each other’s properties, thereby sculpting the evolutionary landscape for both lipids and nucleotides on the prebiotic Earth.

### Experimental Section

#### Materials

Oleic acid (cis-9, C18H34O2, 282.47 g/mol), and glyceryl monooleate (cis-9, C21H40O4, 356.547 g/mol) were purchased from Nu-Check Prep (Elysian, MN, USA) and used without further purification. The rest of the chemicals were purchased from Sigma-Aldrich (Bengaluru, India) and used without further purification. Experiments in buffered conditions were performed in ultrapure water, purified via Millipore (Bedford, MA) Milli-Q system, and used throughout (Resistivity at 25°C = 18 MΩ•cm). Water samples from prebiotically relevant analogue sites were collected during NASA Spaceward Bound expedition to Ladakh, from three separate geothermal pools, i.e., Puga, Chumatang and Panamic, and were used after filtration through a 200 nm Whatman syringe filter.

## Methods

### Preparation of lipid suspensions

The lipid suspensions were prepared by hydrating a thin film of appropriate lipid composition with either 200 mM bicine at pH 8.5 or water samples from the geothermal pools of Puga, Chumatang and Panamic. The pH of water samples from these analogue sites was measured by Joshi et al. in 2017; Puga = 8.48, Chumatang = 8.64, Panamic = 8.37.^[22]^ The thin films were prepared by dissolving appropriate amounts of lipids (only oleic acid or oleic acid + glyceryl monooleate in 2:1 ratio) in a small volume of chloroform, which was then dried under a high vacuum for 4 hours. Buffer/geothermal pool samples and lipid films were heated to 60°C prior to hydration and were equilibrated at the same temperature for 45 minutes post-hydration, during which they were continuously shaken and vortexed to ensure complete solubilization. In the suspensions containing co-solutes, these were added subsequent to the hydration step and equilibration. The concentrations of co-solutes were adjusted to 40 mM of NaCl to the NaCl control (Cont_N_), and 20 mM of nucleotides (A/U/G/C) to the test samples. The lipid suspension was then equilibrated for another 45 minutes at room temperature following the addition of co-solutes. The lipid concentration was always maintained at 4 mM.

### Wet-dry cycles

Wet-dry cycles were set up in glass vials after adding an appropriate amount of the equilibrated lipid suspensions (with co-solutes, if any). Wet-dry cycles were conducted on a heating block set at 90°C in an anoxic and chemically inert environment using a gentle flow of N_2_. Each drying cycle lasted for 1 hour, followed by resuspension using filtered Milli-Q water at room temperature. The suspension was vortexed gently to ensure proper mixing. Aliquots were taken out at different time points for analysis. The volume of Milli-Q used for rehydration was always adjusted during this step to compensate for the amount of sample taken out for analysis after particular time points to maintain the concentration of lipids at 4 mM throughout the cycles. Readings were taken before starting the wet-dry cycles (the 0^th^ cycle) and after completing the 1^st^, 2^nd^, 4^th^, and 8^th^ cycles, respectively.

### Microscopy

Samples were imaged using AxioImager A1, Carl Zeiss upright microscope using 40X magnification (N.A. = 0.75) using Differential Interference Contrast (DIC) modes. 10 μL of the sample was placed on the glass slide and covered with an 18×18 mm glass coverslip. The coverslip was sealed, and the sample was imaged soon thereafter.

Vesicle formation and stability were investigated by observing lipids that can self-assemble into different higher-order structures such as oil droplets and vesicles. We used DIC microscopy to distinguish between these different self-assemblies of lipid suspensions, primarily oil droplets and lipid vesicles. The distinction criterion laid out by Joshi et al. was used to differentiate between the different assemblies.^[22]^ In DIC microscopy images, the appearances of oil droplets and lipid vesicles are distinct. The droplets appear as convex-shaped spheroids and show a high contrast to the background (Supplementary Figure S3a). On the other hand, vesicles appear “doughnut-shaped” and show a low contrast to the background (Supplementary Figure S3b).

### Turbidity estimation

The turbidity of the suspensions was measured using optical density measurements at 450 nm. The optical density was measured using a Shimadzu UV-1800 UV-visible spectrophotometer (Shimadzu Scientific Instruments Inc., Columbia, USA). Readings were taken before starting the wet-dry cycles and after completing the 1^st^, 2^nd^, 4^th^, and 8^th^ cycles, respectively.

### Steady-state fluorescence measurements

To understand the impact of the lipid-nucleotide interactions on the self-assembly of lipid vesicles over multiple wet-dry cycles, the solvatochromic properties of nile red, pyrene and laurdan were utilized. Readings were taken before starting the wet-dry cycles and after completing the 1^st^, 2^nd^, 4^th^, and 8^th^ cycles. Typically, lipid aliquots from these time points were taken out from the main lipid suspensions. These aliquots were then subdivided into four sub-stocks, and either nile red, pyrene or laurdan were added to each sub-stock, and one was left as a blank (sample containing no fluorophore). The concentration of the solvatochromic probes in the suspension was 4 μM and the ratio of the lipid: probe is kept to 1000:1. Lipid suspensions were mixed well and then equilibrated for 20 minutes after the addition of fluorophores. Fluorescence emission was measured on FluoroMax-4C (Horiba Instruments Inc., USA) with a 150 W Xenon, continuous output Ozone-free lamp. The excitation and emission slit-widths were kept at 2 nm. To account for any fluctuations in the intensity resulting from diffractions, emission spectra of blank samples (without any fluorophore) were also obtained. Spectra from the blank samples were subsequently subtracted from the emission of the test samples (containing fluorophores) to obtain the final intensity.

#### Pyrene

The samples containing pyrene were excited at 335 nm, and the emission spectrum was obtained between 350 nm and 600 nm. The ratio of intensities at wavelength 372 nm (I1) and 383 nm (I3) I_1_/I_3_ was taken. The I_1_/I_3_ ratio has been found to be directly correlated with the polarity of Pyrene’s microenvironment.

#### Nile red

Suspensions containing nile red were excited at 530 nm, and the emission spectra were collected between 550 nm and 750 nm. The ratio of intensities at wavelengths 610 nm and 660 nm was used to calculate I_610_/I_660_. The ratio I_610_/I_660_ is inversely correlated with the polarity of the nile red’s microenvironment.

#### Laurdan

The liposomes containing laurdan were excited at 370 nm, and the emission spectrum was collected between wavelengths 400 nm to 600 nm. The generalized polarization (GP) was defined as per equation (1), using intensities at wavelengths 430 nm and 500 nm. GP represents and is directly correlated to the structural order of lipids in a bilayer.

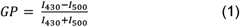

#### Fluorescence anisotropy

Single point anisotropy of laurdan was measured with excitation wavelength 370 nm and emission wavelength 490 nm. Slit-width was kept to 3 nm. Fluorescence anisotropy of laurdan is representative of the fluidity of the bilayer, where the higher the anisotropy, the lower the fluidity. The G factor was automatically measured and accounted for by the instrument before each anisotropy reading.

### Vesicle extrusion

For experiments requiring unilamellar vesicles with narrow size distribution (∼100 nm), suspensions were extruded to achieve the same. Extrusion was performed with an Avanti mini-extruder (Avanti polar lipids, Alabaster, AL, USA) by passing suspensions through a 0.1 μm hydrophilic membrane (Whatman, Buckinghamshire, UK). The suspensions were passed 13 times through the extruder to ensure efficient extrusion at room temperature. Vesicles were stored at 4°C and used within 24 hours of extrusion.

### Calculation of critical aggregation concentration

Critical aggregation concentration (CAC) was estimated using the changes in the emission intensity of 1,6-diphenyl-1,3,5-hexatriene (DPH). The protocol was derived from the earlier work of Sarkar et al.^[37]^ Lipid suspensions of various concentrations were prepared by diluting a lipid stock solution with an oleic acid concentration of 4 mM with 200 mM bicine at pH 8.5. The dilution concentrations were selected to be in the range of previously reported CAC value of OA.^[37]^ Samples were sonicated to ensure the formation of unilamellar vesicles. Co-solutes (nucleotides, NaCl) were added to the suspension after sonication, and samples were equilibrated for 45 minutes at room temperature to ensure proper mixing. Methanol stock of DPH was pipetted into the vial to adjust the final concentration of 2 μM and incubated in the dark for 30 minutes, mixing at 700 RPM, with the temperature adjusted to 40°C. Fluorescence intensity was measured on Tecan Infinite M200 Pro (Tecan, Reading, UK) in an opaque, black 96 well plate. The excitation wavelength used was 350 nm, while the emission wavelength was set to 452 nm.

The fluorescence emission intensity data was fit to a mathematical model plateau followed by one phase association,^[41]^ explained by equation (2).

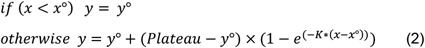

Where, *x*° represents inflexion point (CAC) at which we see rise in intensity (*y*) before a baseline value *y*°, *plateau* represents maximum intensity value, and *K* is the rate constant. Data was plotted and then fitted to this model using graphpad prism 8, and *x*° was extracted from the best fit.^[41]^

### Calculation of normalized differences

For a relative comparison of output values of fluorescence emission and OD between controls and the test systems, as well as across different lipid suspensions, normalized differences of the said property were calculated as per equation (3), for a property X, where subscripts annotate the system. Test systems contain 20 mM nucleotides as co-solutes, while Cont_N_ is a control system containing 40 mM NaCl as a co-solute.

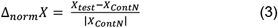

Δ_norm_X is therefore directly proportional to the difference in the value X between the test system and the control system

Error baes for standard error (S.E.) for ∆_norm_X was calculated as per equation (3)

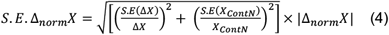

### Statistical analysis

Statistical analysis was used to discern the significance of changes in the analyzed properties (e.g., micropolarity etc.)

#### ANOVA

Two-way repeated-measures ANOVA was conducted using GraphPad Prism 5. The test system containing 20 mM nucleotides as co-solutes were compared to a control system, Cont_N_, that contains 40 mM NaCl. The datasets over the eight wet-dry cycles and four experimental replicates were compared with one another. The changes were considered statistically significant only for ANOVA p-values <0.05.

#### T-test

Two-tailed t-tests were conducted using Microsoft Excel 2016, comparing the 0th cycle with the N^th^ cycle to understand the significance of the wet-dry cycles on the observed properties of the lipid vesicles. The changes were considered statistically significant only for a t-test p-value <0.05.

## Acknowledgements

We acknowledge the microscopy facility at IISER Pune for their support and Prof. Thomas Pucadyil for access to the plate reader. We are grateful to Dr. Ramana Athreya and Dr. Deepak Barua for their helpful discussions on data analysis. We thank Shivani Bodas for her helpful discussions with data fitting in Fig 2. We also thank the COoL lab members, especially Gauri Patki for amazing discussions and useful inputs. KD thanks Kishor Vaigyanik Protsahan Yojana (KVPY) for fellowship. This research was supported by grants from the Science and Engineering Research Board (SERB), Department of Science and Technology, Govt. of India [CRG/2021/001851] and IISER Pune.

