## Supplementary Information for "Nucleotide-protocell interactions: A reciprocal relationship in prebiotically pertinent environments"

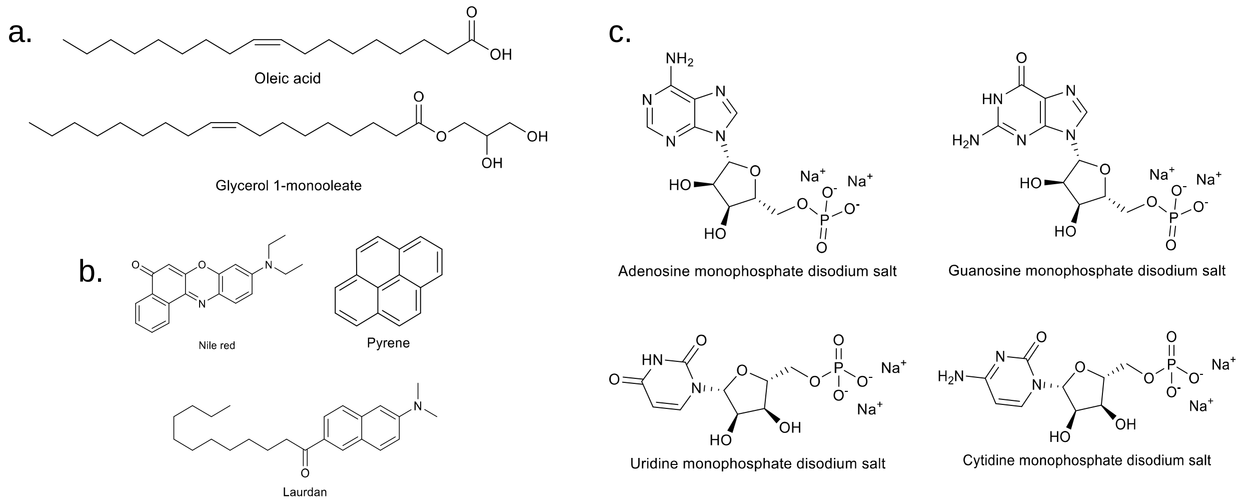


**Figure S1**. Molecular structures of a) amphiphiles (OA, GMO), b) solvatochromic fluorophores (nile red, pyrene and laurdan), and c) disodiated salts of canonical ribose 5'- monophosphates (nucleotides). Structures were drawn using PerkinElmer ChemDraw Professional 20.1.1.

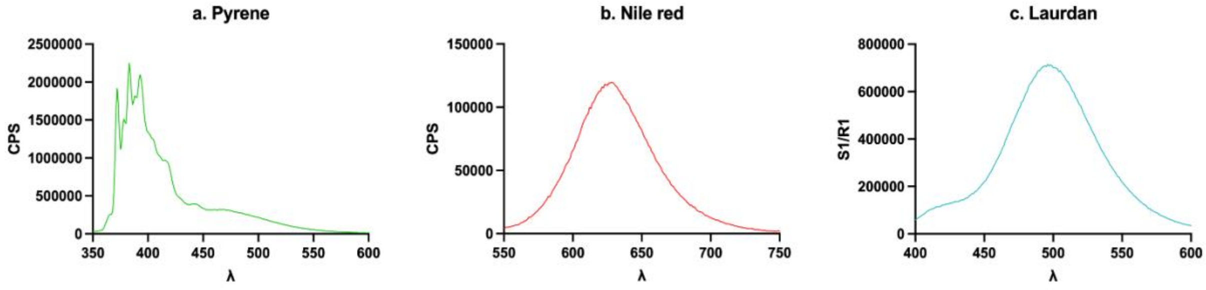


**Figure S2.** Representative emission spectra of the fluorophores used in the study. a) Pyrene: $\lambda_{ex}$ = 335 nm; emission spectrum collected between 350 nm- 600 nm. b) Nile red: $\lambda_{ex}$ = 530 nm; emission spectrum collected between 550 nm- 750 nm. c) Laurdan: $\lambda_{ex}$ = 370 nm; emission spectrum collected between 400 nm- 600 nm.

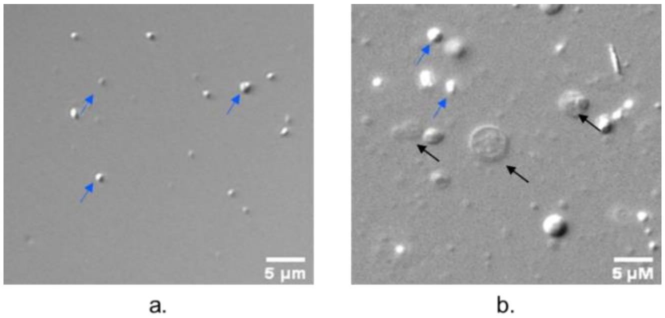


**Figure S3.** Visual distinction between self-assembled structures of lipids. a) Only oil droplets, b) Coexistence of oil droplets and vesicles. Blue and black arrows indicate oil droplets and vesicles respectively. Scale bar is 5 µm. For further details on the distinction protocol, please refer Joshi et al 2017.[1]

**Design of the control experiments**

Control experiments were designed to understand the inherent characteristics of the fatty acid membranes and negate any effect other than that of the interactions with the disodiated salts of nucleotides (NMPdss). To this end, we designed two control experiments. The first set was aimed to discern the effect of wet-dry cycles on the physicochemical properties of membranes in the absence of any co-solutes, which we termed control. The second set was designed to negate the effect of osmotic imbalances arising due to the external addition of co-solutes to the suspensions, which we termed $Cont_N$.

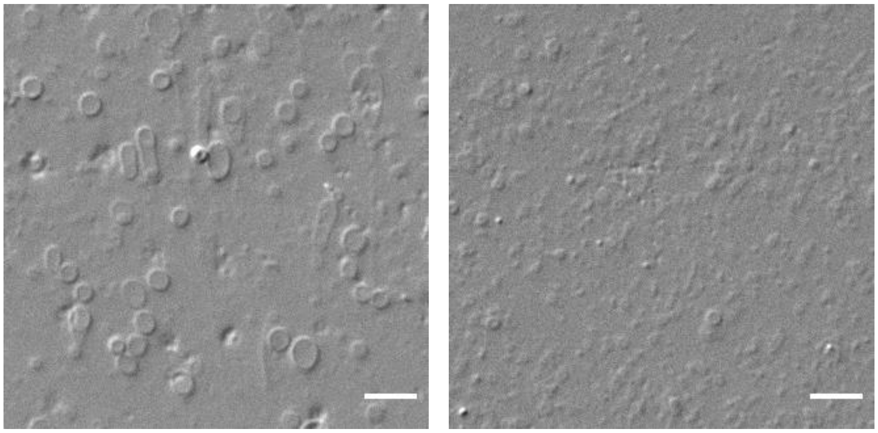


**Figure S4.** Effect of osmotic imbalances on vesicle size due to co-solute addition. The average size of 4 mM OA vesicles (left panel) decreased after the addition and equilibration of 20 mM AMPdss into the OA suspension (right panel). Images were obtained using DIC microscopy at 40x magnification, and the scale bar is 10 µm.

a) **Control**: Control aimed to discern the impact of wet-dry cycles on the properties of model protocell membranes in the absence of any co-solutes. Control entailed vesicles rehydrated only with the dispersion medium, i.e., either 200 mM bicine for experiments conducted in buffered conditions, or with water samples from terrestrial geothermal pools for experiments simulating astrobiologically relevant analogous conditions.

b) **$Cont_N$**: The variety and concentration of cations in the system have been known to affect the stability of fatty acid vesicles.[2,3] Hence, a second set of control experiments termed $Cont_N$ was designed to account for the osmotic imbalances and introduction of Na+ cations to the suspensions due to the addition of NMPdss as co-solutes. $Cont_N$ was prepared by rehydration of fatty acid membranes with dispersion media as mentioned earlier. This was followed by the addition of NaCl stock to adjust the NaCl concentration to 40 mM in the suspensions. This was done to match the sodium concentration in the systems containing NMPdss as co-solutes. For suspensions containing co-solutes, they were added after the hydration of the lipid films. Initial microscopic observation pointed to a qualitative change in the average size of the vesicles after the addition of co-solutes (Figure 3.1, right panel). We reasoned that the change is likely due to osmotic imbalance arising as a consequence of the addition of co-solutes, especially before the start of the wet-dry cycles. The effect of the nucleotide-membrane interactions on the physicochemical properties of model membranes were compared with this $Cont_N$ sample.

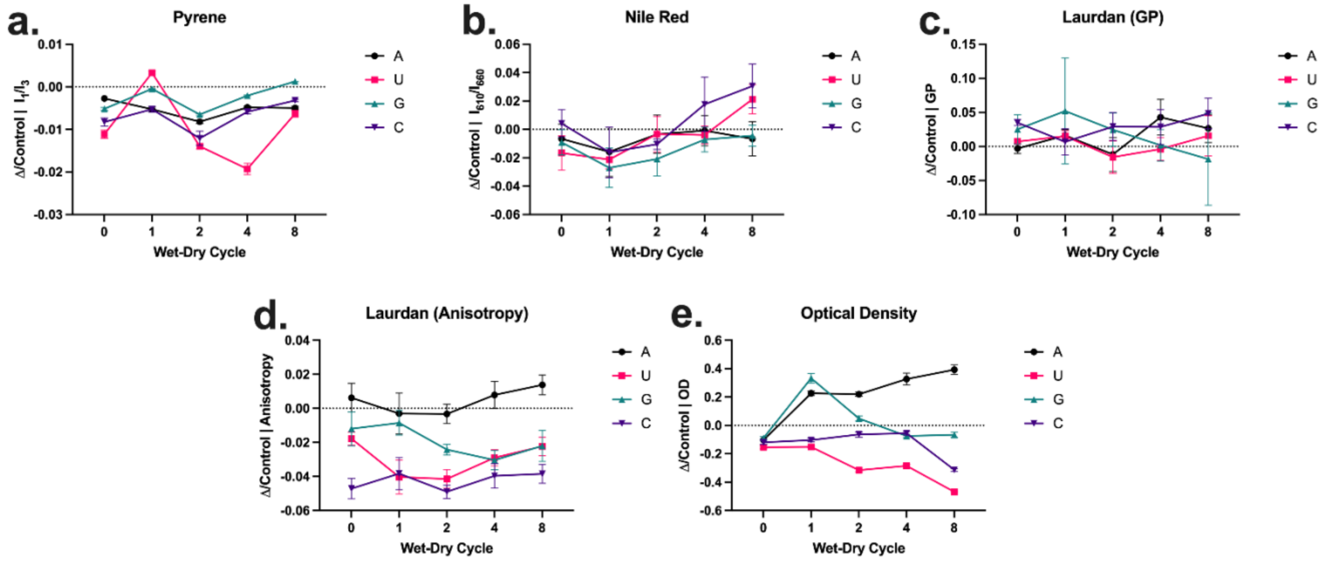


**Figure S5.** Nucleotides affect the physicochemical properties of binary OA-GMO (2:1) membranes in laboratory buffered conditions. The scatter plots represent normalized differences calculated against $Cont_N$ for a) $I_1/I_3$ of pyrene, b) $I_{610}/I_{660}$ of nile red, c) GP of laurdan, d) fluorescence anisotropy of laurdan, e) optical density to understand the effect of nucleotides on protocellular membranes over multiple wet-dry cycles. Timepoints have been appropriately indicated at the x-axis of each data set (as 0-8), and the nucleotide type has been indicated on the right. N = 4, error bars = std. error.

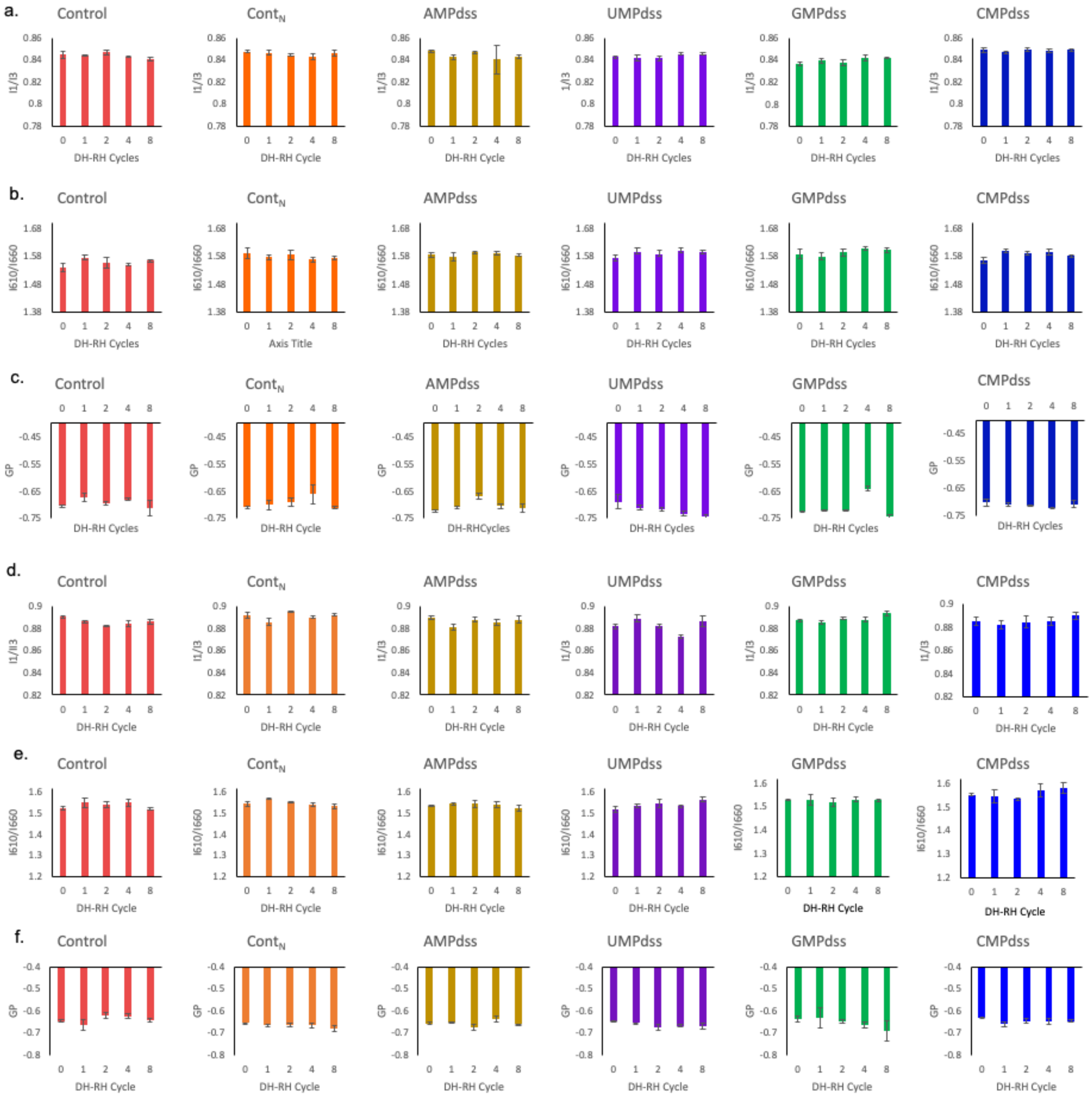


**Figure S6**. Wet-dry cycles do not affect micropolarity and the molecular order of lipids for pure OA and OA-GMO (2:1) membranes in the presence of co-solutes. Data has been represented for a) $I_1/I_3$ of pyrene in OA membranes, b) $I_{610}/I_{660}$ of nile red OA membranes, c) GP of laurdan OA membranes, d) $I_1/I_3$ of pyrene in OA-GMO membranes, e) $I_{610}/I_{660}$ of nile red OA-GMO membranes, f) GP of laurdan OA-GMO membranes. Timepoints have been appropriately indicated on the x-axis of each data set and the nucleotide type involved has been indicated at the top. Control reactions do not have nucleotides. Wet-dry cycles were found largely not to affect these physicochemical properties. N = 4, error bars = std error.

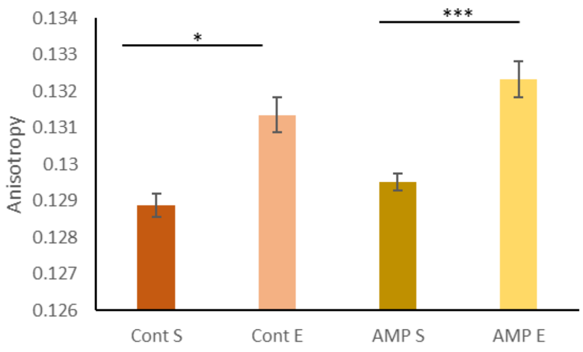


**Figure S7.** Effect of vesicle extrusion on laurdan anisotropy of oleic acid vesicles. 4 mM OA vesicle suspension from rehydration of lipid films were either used as it is (systems indicated with the suffix S), or after extrusion with a membrane of pore size 100 nm (systems indicated with the suffix E). Anisotropy of extruded vesicles was significantly higher than the non-extruded systems. N = 3, error bars = std error. Order of magnitude of statistically significant p-values for the comparative t-test has been represented above the histograms, such that * = 0.01 and *** = 0.0001.

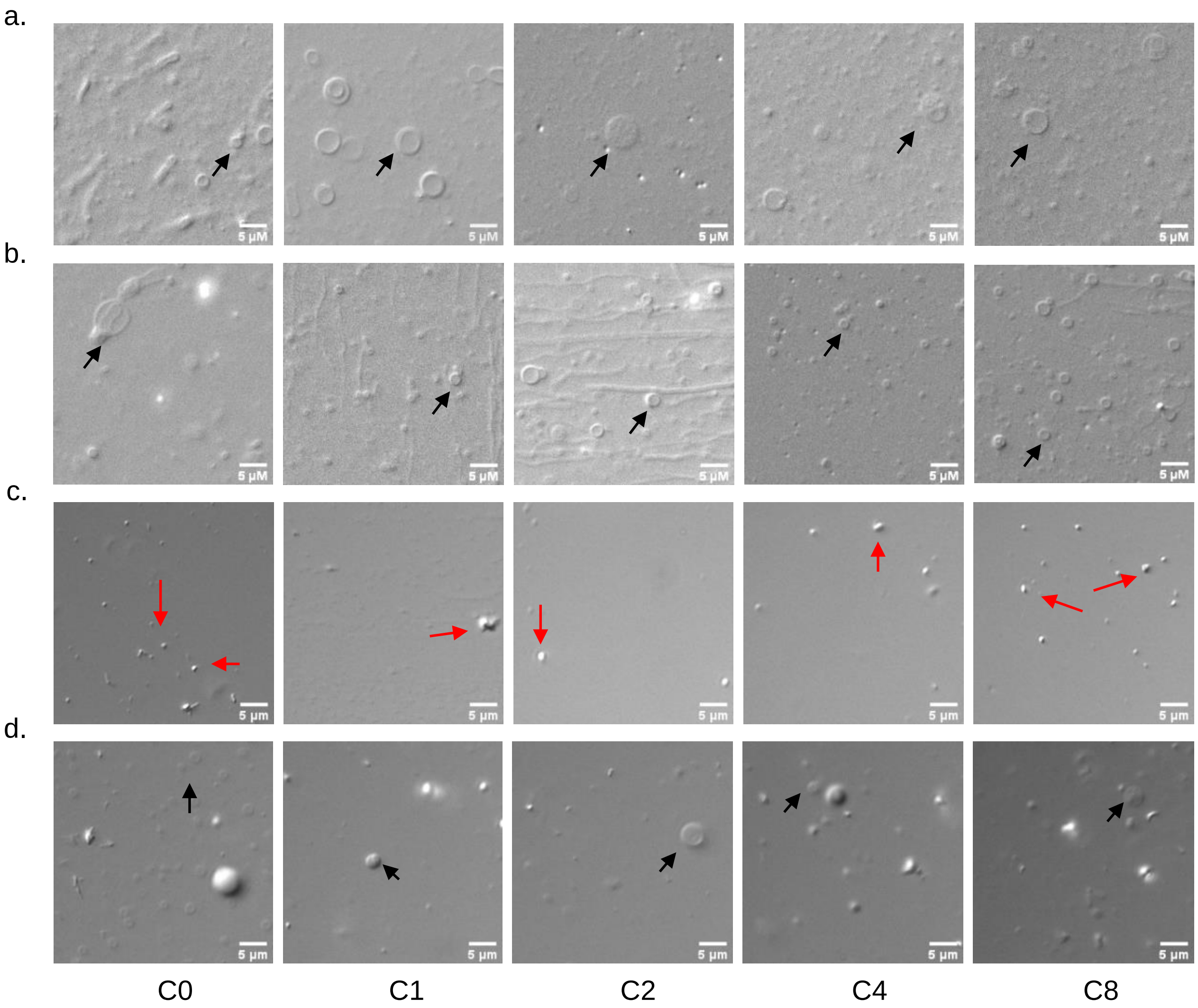


**Figure S8.** Vesicle formation and stability of homogenous and binary systems of OA and OA-GMO over wet-dry cycles under Astrobiologically relevant analogue conditions versus buffered conditions. a) Pure OA suspension in 200 mM bicine buffer, b) OA-GMO (2:1) in 200 mM bicine buffer, c) Pure OA in water sample from Puga, d) OA-GMO (2:1) in water sample from Puga. Pure OA did not form vesicles in the geothermal pool-like conditions, while OA-GMO formed vesicles in both the media. Number of wet-dry cycles are indicated in the bottom panel as C0-C8. Samples were observed using DIC microscopy at 40x, scale bar is 5 µm, N=3. Black arrow= vesicles, red arrow= oil droplets**.**

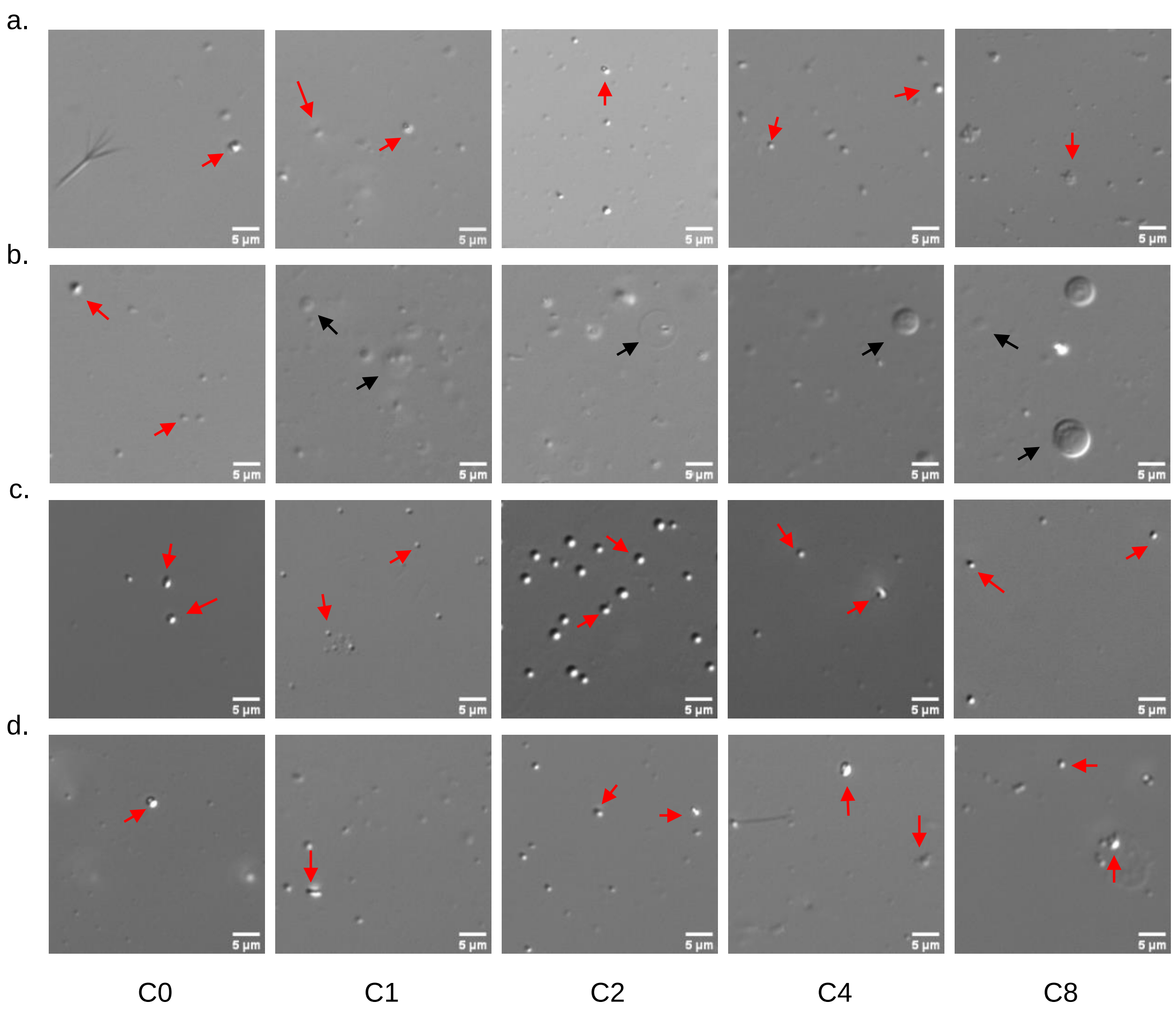


**Figure S9.** Vesicle formation and stability of homogenous and binary systems of OA and OA-GMO over wet-dry cycles without any co-solute under Astrobiologically relevant analogue conditions. a) Pure OA suspension in water sample collected from Chumatang, b) OA-GMO (2:1) in water sample collected from Chumatang, c) Pure OA in water sample from Panamic, d) OA-GMO (2:1) in water sample from Panamic. Pure OA does not form vesicles in geothermal pool-like conditions, while OA-GMO forms vesicles in water sample from Chumatang but not Panamic. Number of wet-dry cycles are indicated in the bottom panel as C0-C8. Samples were observed using DIC microscopy at 40x, scale bar is 5 µm, N=3. Black arrow= vesicles, red arrow= oil droplets.

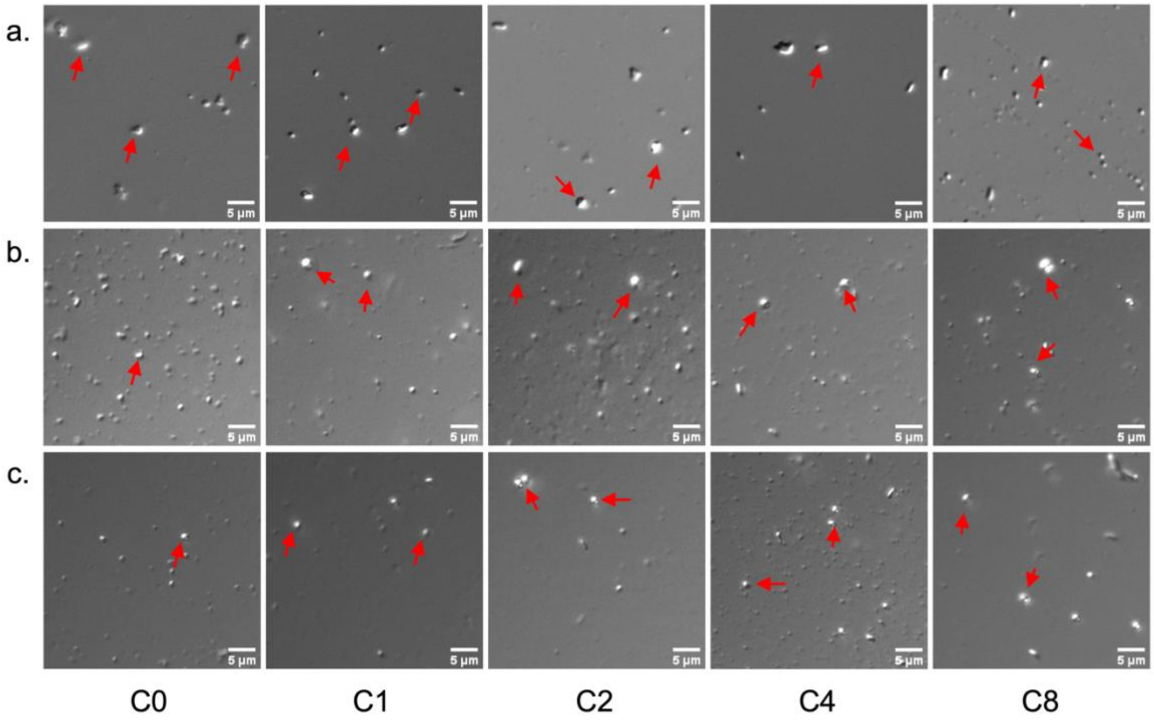


**Figure S10.** Effect of GMP as co-solutes on pure OA vesicle formation and stability in prebiotic analogue conditions. a) Puga + GMP, b) Chumatang + GMP, c) Panamic + GMP We do not observe any vesicle formation in any of the wet-dry cycles. Timepoints have been indicated in the bottom panel as C0-C8. Samples observed using DIC microscopy at 40x, scale bar is 5 µm, N=2. Black arrow= vesicles, red arrow= oil droplets.

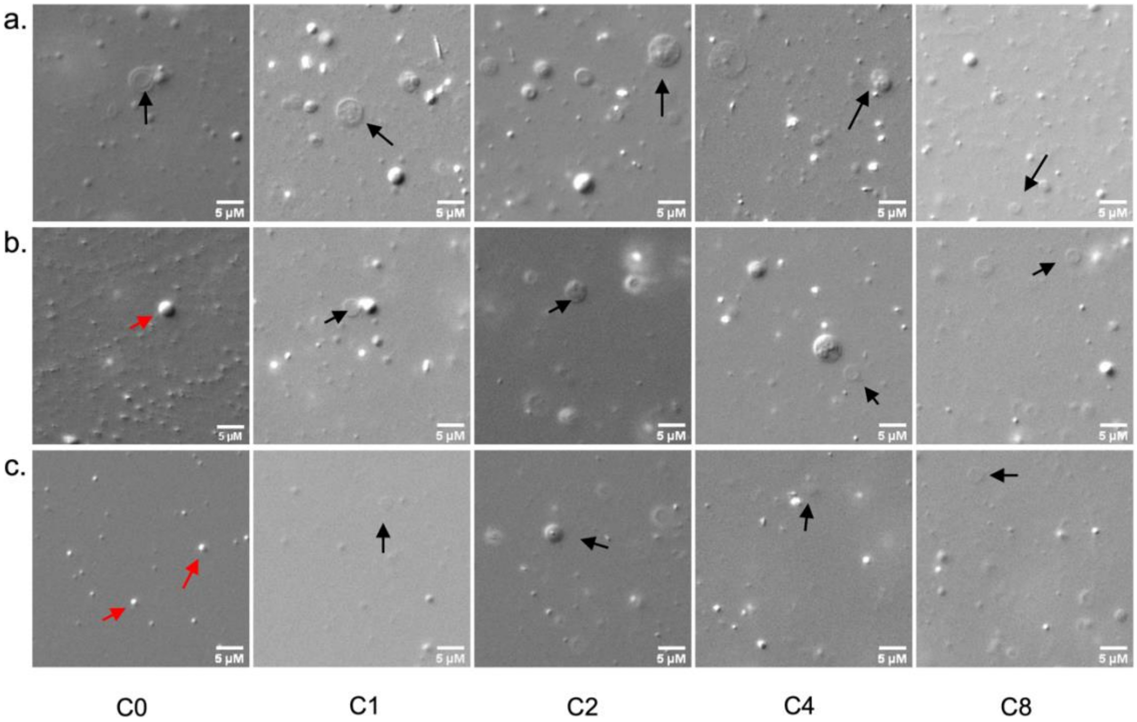


**Figure S11.** Effect of CMP on vesicle formation and stability of binary systems of OA-GMO (2:1) over wet-dry cycles under Astrobiologically relevant analogue conditions. a) Puga + CMP, b) Chumatang + CMP, c) Panamic + CMP. OA-GMO forms vesicles in water sample from all the analogue geothermal pools. Number of wet-dry cycles are indicated in the bottom panel as C0-C8. Samples were observed using DIC microscopy at 40x, scale bar is 5 μm, N=3. Black arrow= vesicles, red arrow= oil droplets**.**

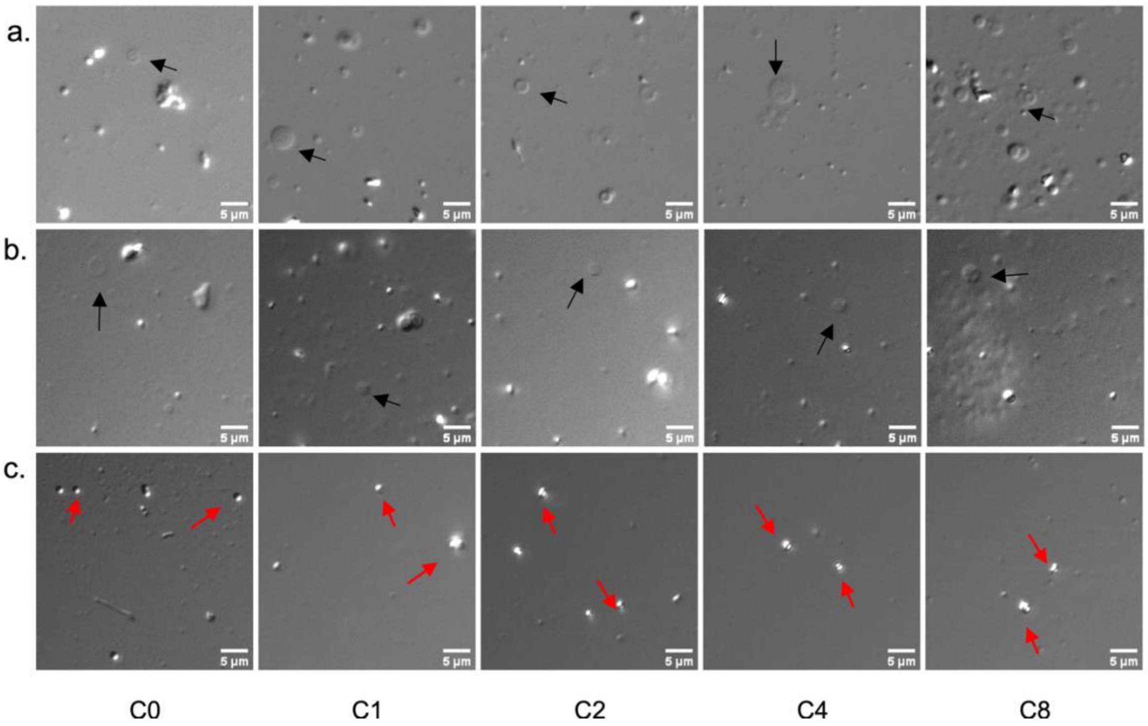


**Figure S12.** Effect of GMP on vesicle formation and stability of binary systems of OA-GMO (2:1) over wet-dry cycles under Astrobiologically relevant analogue conditions. a) Puga + GMP, b) Chumatang + GMP, c) Panamic + GMP. OA-GMO forms vesicles in water samples from Puga and Chumatang, but not in Panamic. Number of wet-dry cycles are indicated in the bottom panel as C0-C8. Samples were observed using DIC microscopy at 40x, scale bar is 5 µm, N=3. Black arrow= vesicles, red arrow= oil droplets**.**

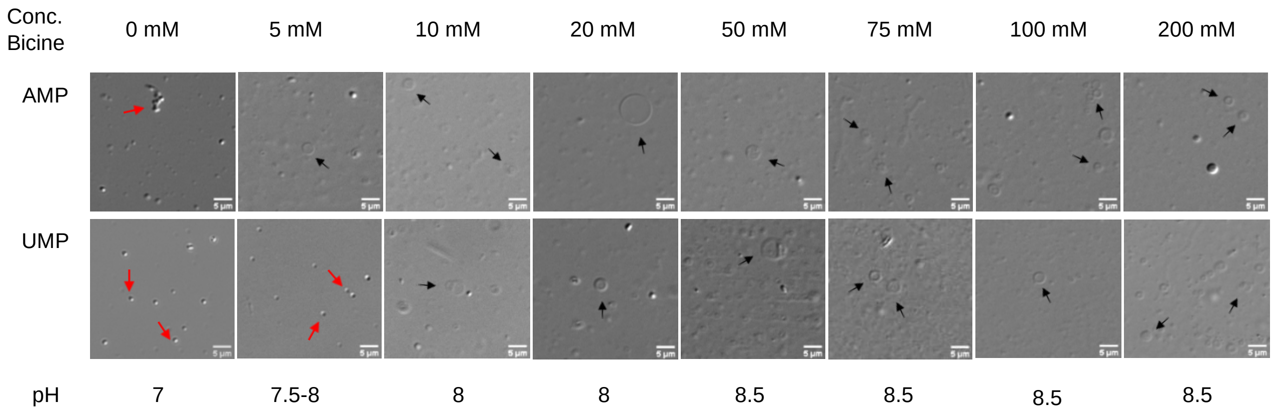


**Figure S13**. Effect of nucleotides on the stability of oleic acid vesicles in different buffer concentrations and across varying pH. Buffer concentration has been indicated in the top panel, while the pH is indicated in the bottom panel. Reactions samples containing OA and UMPdss form vesicles from 200mM till a buffer concentration of 10 mM and a pH of 8. Reactions containing OA and AMPdss form vesicles from 200 mM till a buffer concentration of 5 mM and at a pH between 7.5-8. Samples were observed using DIC microscopy at 40x magnification, scale bar is 5 µm, N=2. Black arrow= vesicles, red arrow= oil droplets. For overall comparison of different systems refer Table ST7

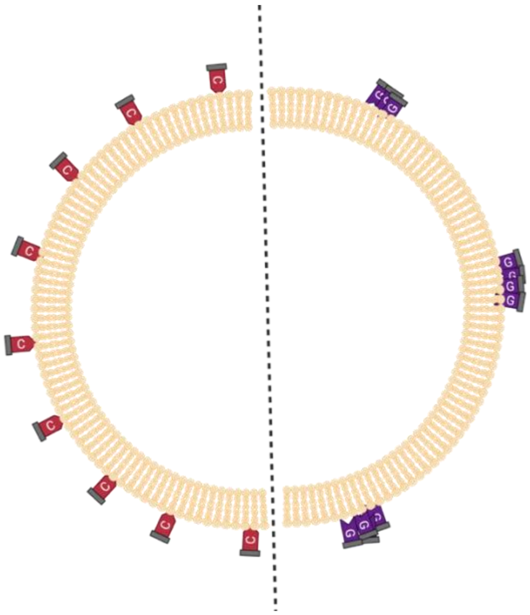

**Figure S14.** Schematic illustration of nucleotide-membrane interactions and the proposed difference between the interaction of CMP and GMP. Nucleobase and ribose interact with the fatty acid membranes, while phosphate points out towards the water. Illustration created with BioRender.com

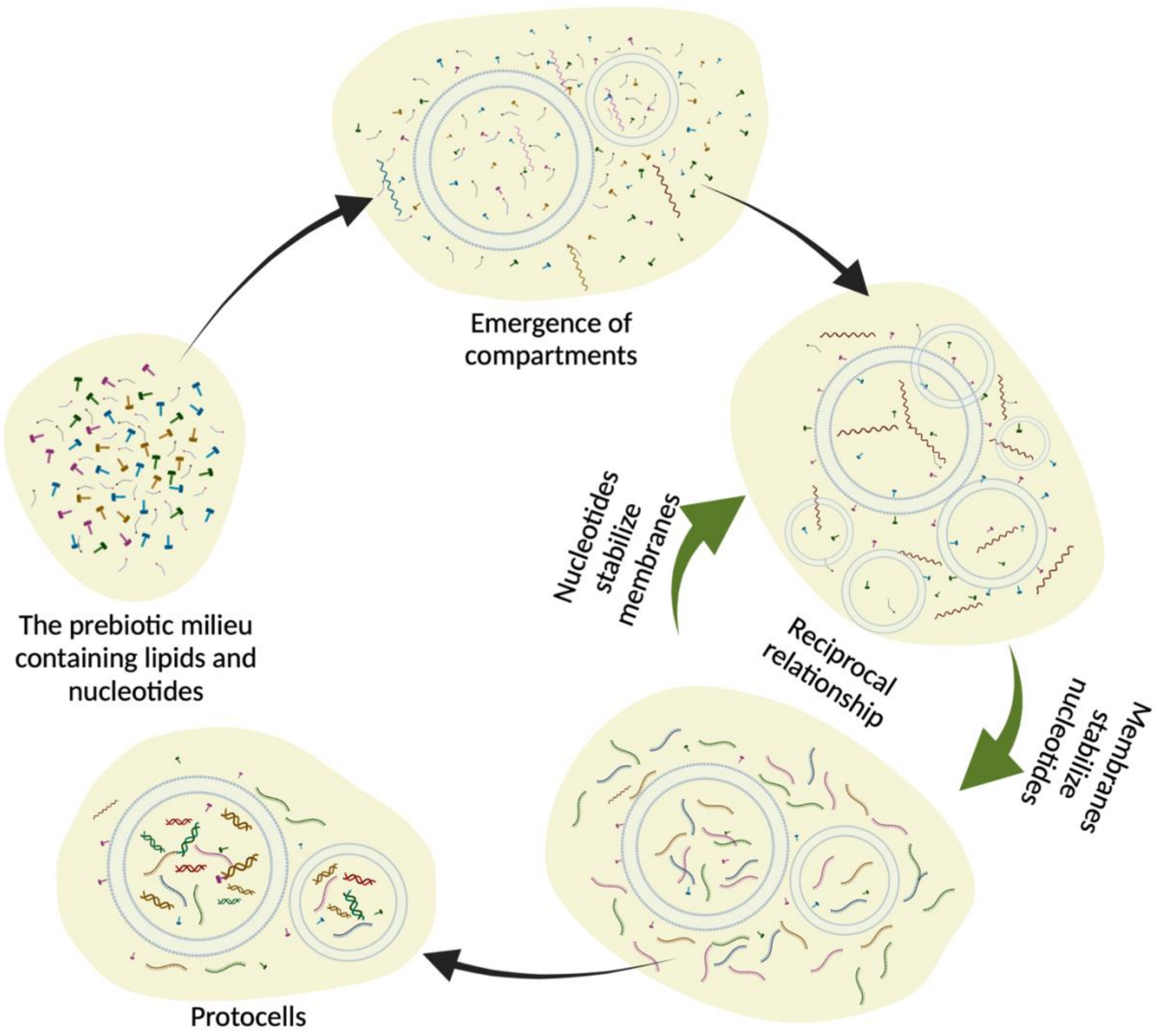


**Figure S15** Schematic illustration of the reciprocal relationship between nucleotides and protocellular membranes. Illustration created with BioRender.com

### Supplementary Tables

| OA-GMO (2:1) | AMP | UMP | GMP | CMP |
|---|---|---|---|---|
| Micropolarity near the headgroup | **0.0271** | **0.004** | 0.1573 | **0.0063** |
| Micropolarity at the center of the bilayer | **0.028** | 0.3351 | **0.0446** | 0.5503 |
| Molecular order of lipids | 0.1545 | 0.7356 | 0.068 | 0.3012 |
| Membrane fluidity | **<0.0001** | **<0.0001** | **0.0008** | **<0.0001** |
| Turbidity | **<0.0001** | **<0.0001** | **0.0397** | **<0.0001** |

**Table ST1.** Comparison of p-values obtained from two-way Repeated Measures ANOVA analysis performed for the test samples and the $Cont_N$ for OA-GMO vesicles. The physicochemical properties were compared for the test systems with $Cont_N$ over eight wet-dry cycles to gain a systems level perspective of the effect of nucleotides on the physicochemical properties of protocellular membranes. p-values smaller than 0.05 were considered statistically significant and highlighted in bold text.

| GMO (2:1) Puga Vs Buffered | $Cont_N$ | GMP | CMP |
|---|---|---|---|
| Micropolarity near the headgroup | **0.0083** | **<0.0001** | **0.0002** |
| Micropolarity at the centre of the bilayer | 0.4854 | **0.0274** | 0.4515 |
| Molecular order of lipids | 0.0562 | 0.0651 | 0.5219 |
| Membrane fluidity | **0.0325** | **0.0103** | **0.0008** |
| Turbidity | **0.0003** | **<0.0001** | **<0.0001** |

**Table ST2.** p-values for two-way Repeated Measures ANOVA for the OA-GMO samples in geothermal pool sample as compared with suspensions prepared in 200 mM bicine. The physicochemical properties of the suspensions prepared in Puga water were compared against their buffered counterparts over eight wet-dry cycles to gain a system level perspective of the effect of ionic diversity of the medium, on the physicochemical properties of protocellular membranes. p-values smaller than 0.05 were considered statistically significant and highlighted in bold text.

| OA | | | | | | |
|---|---|---|---|---|---|---|
| Pyrene | | | | | | |
| Cycle | Control | $Cont_N$ | AMP | UMP | GMP | CMP |
| 0 | ------------- | ------------- | ------------- | ------------- | ------------- | ------------- |
| 1 | 0.7962 | 0.7105 | 0.06 | 0.7856 | 0.3943 | 0.4104 |
| 2 | 0.5787 | 0.1052 | 0.4794 | 0.7285 | 0.694 | 0.8096 |
| 4 | 0.6884 | 0.1748 | 0.2135 | 0.1581 | 0.1372 | 0.7708 |
| 8 | 0.6884 | 0.6417 | 0.0498 | 0.1617 | 0.0308 | 0.9863 |
| Nile red | | | | | | |
| Cycle | Control | $Cont_N$ | AMP | UMP | GMP | CMP |
| 0 | ------------- | ------------- | ------------- | ------------- | ------------- | ------------- |
| 1 | 0.0902 | 0.4702 | 0.6266 | 0.1752 | 0.71 | 0.04921 |
| 2 | 0.4802 | 0.818 | 0.4707 | 0.4919 | 0.8353 | 0.109 |
| 4 | 0.5623 | 0.2733 | 0.7575 | 0.0804 | 0.3729 | 0.0816 |
| 8 | 0.5622 | 0.3659 | 0.8988 | 0.1575 | 0.5186 | 0.254 |
| Laurdan (GP) | | | | | | |
| Cycle | Control | $Cont_N$ | AMP | UMP | GMP | CMP |
| 0 | ------------- | ------------- | ------------- | ------------- | ------------- | ------------- |
| 1 | 0.1457 | 0.6602 | 0.0642 | 0.5048 | 0.4494 | 0.5642 |
| 2 | 0.3048 | 0.3062 | 0.0115 | 0.3798 | 0.4656 | 0.3951 |
| 4 | 0.1322 | 0.2115 | 0.1332 | 0.2127 | 0.0004 | 0.199 |
| 8 | 0.3451 | 0.8182 | 0.5383 | 0.1454 | 0.0665 | 0.7727 |
| Laurdan (Anisotropy) | | | | | | |
| Cycle | Control | $Cont_N$ | AMP | UMP | GMP | CMP |
| 0 | ------------- | ------------- | ------------- | ------------- | ------------- | ------------- |
| 1 | 0.0005 | <0.0001 | 0.0003 | 0.0068 | 0.0063 | <0.0001 |
| 2 | <0.0001 | <0.0001 | <0.0001 | 0.0044 | 0.2111 | 0.0011 |
| 4 | 0.0026 | <0.0001 | <0.0001 | 0.0028 | 0.0233 | 0.0005 |
| 8 | 0.0026 | <0.0001 | 0.0033 | 0.00256 | 0.124 | <0.0001 |
| Turbidity | | | | | | |
| Cycle | Control | $Cont_N$ | AMP | UMP | GMP | CMP |
| 0 | ---------------- | ---------------- | ---------------- | ---------------- | ---------------- | ---------------- |
| 1 | <0.0001 | 0.0002 | 0.0003 | <0.0001 | <0.0001 | 0.0001 |
| 2 | <0.0001 | 0.0002 | 0.0008 | 0.0003 | <0.0001 | 0.0011 |
| 4 | 0.0001 | 0.624 | 0.0008 | 0.0234 | 0.001 | 0.0101 |
| 8 | <0.0001 | 0.0008 | 0.0023 | <0.0001 | <0.0001 | 0.0011 |

**Table ST3.** p-values for two-tailed t-tests comparing $0^{th}$ cycle with $N^{th}$ cycle for OA suspensions to investigate the statistical significance of change in the physicochemical properties of protocellular membranes due to wet-dry cycles. p-values smaller than 0.05 were considered statistically significant.

| OA-GMO (2:1) | | | | | | |
|---|---|---|---|---|---|---|
| Pyrene | | | | | | |
| Cycle | Control | $Cont_N$ | AMPdss | UMPdss | GMPdss | CMPdss |
| 0 | ------------- | ------------- | ------------- | ------------- | ------------- | ------------- |
| 1 | 0.0272 | 0.1635 | 0.0355 | 0.1197 | 0.48 | 0.5568 |
| 2 | 0.0022 | 0.2613 | 0.5494 | 0.8267 | 0.2757 | 0.966 |
| 4 | 0.1292 | 0.4881 | 0.2539 | 0.0139 | 0.7303 | 0.9616 |
| 8 | 0.1292 | 0.4881 | 0.7312 | 0.4129 | 0.0298 | 0.338 |
| Nile red | | | | | | |
| Cycle | Control | $Cont_N$ | AMPdss | UMPdss | GMPdss | CMPdss |
| 0 | ------------- | ------------- | ------------- | ------------- | ------------- | ------------- |
| 1 | 0.3194 | 0.1615 | 0.2499 | 0.3078 | 0.8957 | 0.8265 |
| 2 | 0.3453 | 0.7099 | 0.5796 | 0.3018 | 0.5418 | 0.0733 |
| 4 | 0.2174 | 0.8337 | 0.723 | 0.3008 | 0.9727 | 0.5887 |
| 8 | 0.2174 | 0.8337 | 0.4697 | 0.0338 | 0.5886 | 0.2672 |
| Laurdan (GP) | | | | | | |
| Cycle | Control | $Cont_N$ | AMPdss | UMPdss | GMPdss | CMPdss |
| 0 | ------------- | ------------- | ------------- | ------------- | ------------- | ------------- |
| 1 | 0.5338 | 0.1151 | 0.7007 | 0.1675 | 0.765 | 0.0933 |
| 2 | 0.1875 | 0.9999 | 0.3296 | 0.1432 | 0.2432 | 0.2776 |
| 4 | 0.8716 | 0.9966 | 0.2581 | 0.0204 | 0.0251 | 0.3777 |
| 8 | 0.1816 | 0.4483 | 0.4404 | 0.2971 | 0.0959 | 0.1121 |
| Laurdan (Anisotropy) | | | | | | |
| Cycle | Control | $Cont_N$ | AMPdss | UMPdss | GMPdss | CMPdss |
| 0 | ------------- | ------------- | ------------- | ------------- | ------------- | ------------- |
| 1 | 0.134 | 0.9384 | 0.35 | 0.0501 | 0.7919 | 0.386 |
| 2 | 0.4248 | 0.0656 | 0.0703 | 0.0039 | 0.1482 | 0.2634 |
| 4 | 0.0007 | 0.0007 | 0.0559 | 0.0017 | 0.0246 | 0.2876 |
| 8 | 0.0007 | 0.0007 | 0.0059 | 0.0016 | 0.0319 | 0.0601 |
| Turbidity | | | | | | |
| Cycle | Control | $Cont_N$ | AMPdss | UMPdss | GMPdss | CMPdss |
| 0 | ---------------- | ---------------- | ---------------- | ---------------- | ---------------- | ---------------- |
| 1 | 0.0025 | 0.0001 | <0.0001 | 0.0011 | 0.0065 | 0.0183 |
| 2 | 0.3068 | 0.0016 | <0.0001 | 0.0003 | 0.0086 | 0.0008 |
| 4 | 0.0181 | 0.0257 | 0.0019 | 0.0001 | 0.0147 | 0.0745 |
| 8 | <0.0001 | 0.0002 | 0.0007 | <0.0001 | 0.0001 | <0.0001 |

**Table ST4.** p-values for two-tailed t-tests comparing $0^{th}$ cycle with $N^{th}$ cycle for OA-GMO (2:1) suspensions to investigate the statistical significance of change in the physicochemical properties of protocellular membranes due to wet-dry cycles. p-values smaller than 0.05 were considered statistically significant.

| | A | B | C | D | E | F |
|---|---|---|---|---|---|---|
| Buffer Conc. | Buffer | Buffer + OA | B + AMP | B + UMP | B + GMP | B + CMP |
| 0 mM | 7 | 6.5 | 7 | 7 | 7 | 7 |
| 5 mM | 7.5-8 | 7.5-8 | 7.5-8 | 7.5-8 | 7.5-8 | 7.5-8 |
| 10 mM | 8 | 8 | 8 | 8 | 8 | 8 |
| 20 mM | 8 | 8-8.5 | 8-8.5 | 8-8.5 | 8 | 8 |
| 50 mM | 8.5 | 8.5 | 8.5 | 8.5 | 8.5 | 8.5 |
| 75 mM | 8.5 | 8.5 | 8.5 | 8.5 | 8.5 | 8.5 |
| 100 mM | 8.5 | 8.5 | 8.5 | 8.5 | 8.5 | 8.5 |
| 200 mM | 8.5 | 8.5 | 8.5 | 8.5 | 8.5 | 8.5 |

**Table ST5**. pH of OA suspensions made using different bicine buffer dilutions.

| | A | B | C | D | E |
|---|---|---|---|---|---|
| Buffer Conc. | Buffer + OA | A + AMP | A+ UMP | A+ GMP | A+ CMP |
| 0 mM | ✗ | ✗ | ✗ | ✗ | ✗ |
| 5 mM | ✗ | ✓ | ✗ | ✗ | ✓ |
| 10 mM | ✗ | ✓ | ✓ | ✗ | ✓ |
| 20 mM | ✗ | ✓ | ✓ | ✗ | ✓ |
| 50 mM | ✓ | ✓ | ✓ | ✓ | ✓ |
| 75 mM | ✓ | ✓ | ✓ | ✓ | ✓ |
| 100 mM | ✓ | ✓ | ✓ | ✓ | ✓ |
| 200 mM | ✓ | ✓ | ✓ | ✓ | ✓ |

**Table ST6**. Formation of oleic acid vesicles under varying bicine concentration and pH. Lipid films were rehydrated with different concentrations of the bicine buffer. The propensity of oleic acid to form vesicles under these conditions was studied using DIC microscopy in the presence and absence of nucleotides. ✓ = vesicles observed, ✗= no vesicles observed.
